# Model-driven engineering of *Cutaneotrichosporon oleaginosus* ATCC 20509 for improved microbial oil production

**DOI:** 10.1101/2024.03.19.585731

**Authors:** Zeynep Efsun Duman-Özdamar, Mattijs K. Julsing, Janine A.C. Verbokkem, Emil Wolbert, Vitor A.P. Martins dos Santos, Jeroen Hugenholtz, Maria Suarez-Diez

## Abstract

Consumption of plant-based oils, especially palm oil, is increasing at an alarming rate. This boosted demand for palm oil has drastic effects on the ecosystem as its production is not sustainable. *C. oleaginosus* is an oleaginous yeast with great potential as a source for microbial-based oil production which is a sustainable alternative to palm oil. However, microbial processes are not yet economically feasible to replace palm oil, unto a large extent due to limited lipid accumulation in the microbe, which limits titers and productivity. Therefore, obtaining enhanced lipid accumulation is essential to render this process commercially viable. Herein we deployed a systematic, iterative Design-Build-Test-Learn (DBTL) approach to establish *C. oleaginosus* as an efficient fatty acid production platform. In the design step, we identified genes and medium supplements that improved lipid content. To this end, we compared its transcriptional landscape in conditions with high and low amounts of lipid production. A metabolic map was reconstructed and integrated with the expression data. Finally, the genome-scale metabolic model of *C. oleaginosus* was used to explore metabolism under maximal growth and maximal production conditions. The combination of these four analyses led to the selection of four overexpression targets (ATP-citrate lyase (*ACL1*), acetyl-CoA carboxylase (*ACC*), threonine synthase (*TS*), and hydroxymethylglutaryl-CoA synthase (*HMGS*)) and five media supplements (biotin, thiamine, threonine, serine, and aspartate). We established an electroporation-based co-transformation method to implement selected genetic interventions. These findings were experimentally validated in the build and test steps of the DBTL approach by adding supplements into the medium and overexpressing the identified genes. Characterization of ACL, ACC, and TS at various C/N ratios, and the addition of medium supplements provided up to 56% (w/w) lipid content, and a 2.5-fold increase in total lipid in the glycerol and urea-based defined medium. In the learn step, quadratic models identified the optimum C/N ratios shifted towards around C/N240. These results firmly confirm *C. oleaginous* as a sustainable alternative to replace palm as an oil source.

**Highlights:**

- Transcriptional profile and metabolic model analyzed, predicting genetic targets and medium supplements.
- Genetic targets and medium supplements for improved oil production.
- The genetic toolbox for *C. oleaginosus* was expanded (co-transformation method, promoters, genes, and terminators).
- Experimental validations showed that biotin, and threonine increased lipid content.
- Overexpression of *ACL1, ACC,* and *TS* in *C. oleaginosus* provided higher oil content.

**Graphical Abstract:** 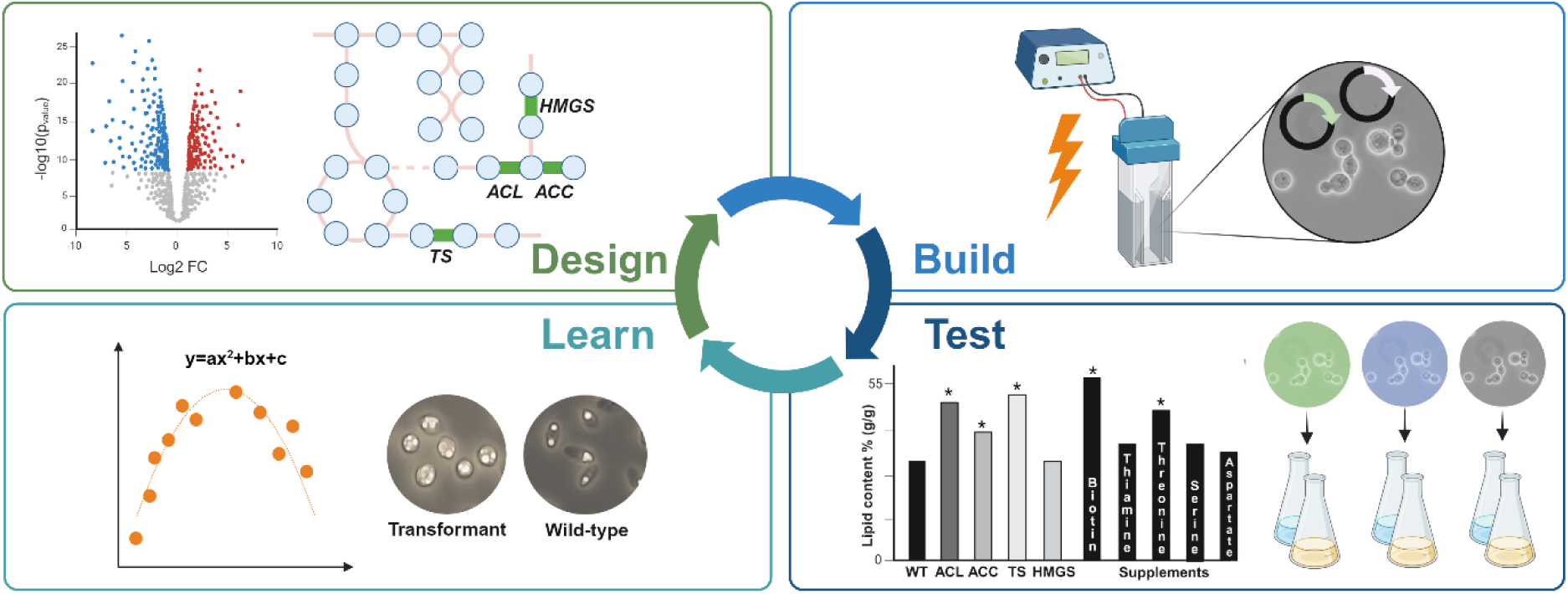

## 1. Introduction

Plant-based oils and fatty acids are widely used in food, feed, chemical, personal care, and cosmetic products as they provide beneficial functions to these products such as texture, flavor or extended shelf-life (Holley and Patel, 2005; P. Desbois, 2012; Rustan and Drevon, 2005). Palm oil is especially preferred due to it being a cheap source of these functional components. The increasing demand for palm oil has led to the replacement of the native tropical forests, and traditional vegetation in many Asian, South American, and African countries. This replacement not only poses a threat to the local ecosystem but also has severe implications for local livelihoods, contributing to deforestation and climate change (Abubakar et al., 2021; Snashall and Poulos, 2021). Although efforts have been implemented to combat deforestation through initiatives like the Roundtable on Sustainable Palm Oil (RSPO), the use of palm oil remains contentious (Cazzolla Gatti et al., 2019). To that end, developing a sustainable alternative to plant-based fatty acids and oils is urgent and of utmost interest.

Oil-producing yeasts, referred to as oleaginous yeasts, have strong potential as sustainable alternatives for lipid production in various industrial applications (Patel et al., 2020). An oleaginous yeast *Cutaneotrichosporon oleaginosus,* (formerly also known as *Apiotrichum curvatum, Cryptococcus curvatus, Trichosporon cutaneum, Trichosporon oleaginosus,* and *Cutaneotrichosporon curvatum*) is able to accumulate oils at contents exceeding 40% of their dry biomass under carefully selected conditions. (Beopoulos et al., 2011; Iassonova et al., 2008). The accumulation of lipids in oleaginous yeasts is induced by limiting specific nutrients such as nitrogen. *C. oleaginosus* starts to accumulate oils by redirecting the excess carbon to be stored as lipids when the critical carbon-to-nitrogen (C/N) ratio of 11 g/g is reached (Carsanba et al., 2018; Ykema et al., 1986). Extensive biochemical and genetic engineering studies revealed that in N-limiting conditions a continuous supply of acetyl-CoA, malonyl-CoA, and NADPH is required for fatty acid biosynthesis in *C. oleaginosus* (Pham et al., 2021; Robles-Iglesias et al., 2023). In addition to the potentially high oil contents, *C. oleaginosus* can use a large variety of carbon sources as substrates including glucose, xylose, cellobiose, sucrose, glycerol, and lactose, and utilize waste streams such as whey permeate, volatile fatty acids, office paper production waste streams, and crude glycerol from biodiesel production (Annamalai et al., 2018; Dobrowolski et al., 2020; Liu et al., 2017). Due to these advantages, *C. oleaginosus* has been flagged as one of the promising microbial oil-producing cell factories.

On the other hand, Koutinas et al. reported that the implementation of microbial oil in industrial applications is not yet economically feasible. For *C. oleaginosus*, it has been calculated that the process would only become economically feasible if lipid accumulation increases (Koutinas et al., 2014). Karamerou et al. suggested using whole cells in a product, optimizing process operation mode, and improving the downstream processing as a complementary approach to achieve a feasible production process when lipid accumulation is around 60% (w/w) (Karamerou et al., 2021).

Recently, efforts have been made to design *C. oleaginosus* as efficient cell factories by using metabolic engineering strategies and optimizing the medium and cultivation conditions for enhancing yield and lipid productivity (Adrio, 2017; Shi and Zhao, 2017). Previously, we extensively investigated the effect of C/N ratio and temperature on the physiology of oleaginous yeasts and showed that the lipid accumulation profile of *C. oleaginosus* was strongly dependent on these factors (Duman-Özdamar et al., 2022). Furthermore, (Awad et al., 2019) reported carbon and nitrogen sources affected the growth and lipid content of *C. oleaginosus*. Implementation of mitochondrial pyruvate dehydrogenase (PDH) bypass into *C. oleaginosus* by overexpression of pyruvate decarboxylase, acetaldehyde dehydrogenase, and acetyl-CoA synthase from *Saccharomyces cerevisiae* besides diacylglycerol acyltransferase from *Rhizopus oryzae* has resulted in an increased triacylglycerol (TAG) yield by using xylose as substrates at an N-limiting minimal medium (Koivuranta et al., 2018). CRISPR-mediated genetic engineering tool was developed and reported by Shaigani et al., 2023 and the Δ9-desaturase gene was overexpressed, and the Δ12-desaturase gene was overexpressed and knocked out to tailor the fatty acid composition.

Despite these developments, there are still limited strain engineering studies on *C. oleaginosus*. Recent advances in systems biology and computational modeling such as constrained-based genome-scale metabolic models (GEM models) have broadened the way of finding new strategies for strain engineering (Machado and Herrgård, 2015) and to engineer oleaginous yeasts that have naturally high lipid accumulation (Shi and Zhao, 2017). The translation from genotype to the desired phenotype is a multi-step process that requires combinatorial work on pathway and metabolism represented by constrained-based GEM models, implementing genetic modifications, and adapting culture conditions. To achieve the full potential of these oleaginous yeasts as lipid producers, a systematic and efficient strategy needs to be followed.

In this study, we followed the Design-Build-Test-Learn (DBTL) approach, which represents a streamlined approach to iterate the steps of strain development by facilitating the systems biology, metabolic engineering, bioprocess engineering and modeling framework, to develop a sustainable and more productive fatty acid production platform using *C. oleaginosus*. To this end, we systematically intertwined predictions from the genome-scale metabolic model and characterization of the transcriptional landscape under differential oil-producing conditions with experimental validations on wild-type and newly constructed transformants.

## 2. Materials and Methods

### 2.1. Analysis of transcriptome data

Three strains of *Cutaneotrichosporon oleaginosus* (wild-type, Δ9, Δ12) were grown in high C/N ratio and low C/N ratio condensed distilled soluble (CDS, crude glycerol) medium. Detailed protocol for experiments performed to generate RNA-seq data can be found in Supplementary File A. RNA-seq datasets are available at ENA study PRJEB73355.

Reads were quality-filtered and trimmed using fastp v0.20.0 (Chen et al., 2018). Ribosomal RNA was removed using bbduk from the BBMap v38.57 using the available ribokmers.fa.gz file in the BBMap suite (Bushnell, 2014). Read inspection was performed using FastQC v0.11.9 and MultiQC v1.8 (Ewels et al., 2016; Simon Andrews, 2020).

The structurally annotated genome of *C. oleaginosus* ATCC20509 was retrieved from ENA study PRJEB73837 (Nijsse et al., 2024). The filtered reads were mapped to the genome and its FUNGAP predicted genes using RSEM v1.3.1 and STAR v2.7.3a (Li and Dewey, 2011).

The package txtimport was used to import the data in R and obtain summary statistics Differential expression analysis was performed using DEseq2 and p-values were corrected for multiple testing using Benjamini-Hochberg(Love et al., 2014; Soneson et al., 2015). For each comparison, genes with a TPM of at least 100 in both replicates in a condition were kept for further investigation. Genes were considered differentially expressed when corrected p-value ≤ 0.05. Downregulated and upregulated genes were additionally filtered based on log_2_FoldChange (−0.58 ≥ log_2_FoldChage ≥ 0.58, corresponding to fold changes of 1.5 times

### 2.2. Gene Ontology (GO) enrichment analysis

Genes were functionally annotated using the EGGNog mapper (Cantalapiedra et al., 2021). Enrichment analysis was performed on differentially expressed genes, up-regulated and down-regulated, by applying a hypergeometric test via the BiNGO Cytoscape app (v 3.7.2) (Maere et al., 2005) and considering the annotated genome as the background set. Benjamini & Hochberg false discovery rate (FDR) correction method was used to control FDR and results were deemed significant when FDR < 0.05. Annotations used in this analysis can be found in supplementary file B.

### 2.3. Pathway visualization

A metabolic map consisting of central carbon metabolism, lipid synthesis pathway, and fatty acid elongation pathway was constructed in Escher (https://escher.github.io/) based on the constrained based genome-scale metabolic model (GEM) of *C. oleaginosus*, *i*NP636_Coleaginosus_ATCC20509_v2 (Rosmalen et al., 2023) which is an updated version of *i*NP636_Coleaginosus_ATCC20509 (Pham et al., 2021). The *i*NP636_Coleaginosus_ATCC20509 was manually curated by adding missing fatty acid elongation reactions (Rosmalen et al., 2023). Afterward, differentially expressed genes were imported into the constructed metabolic map and visualized based on their log2FoldChange values in Escher (supplementary file C).

### 2.4. Comparative flux sampling analysis using the GEM

The *i*NP636_Coleaginosus_ATCC20509_v2 model was used to simulate scenarios of maximum growth, maximum lipid production, and slow growth. In each scenario flux sampling was used to characterize the metabolic space and reactions with the highest changes between scenarios were selected as targets for further inspection as described by (Rosmalen et al., 2023).

### 2.5. Strains, media, and growth conditions in shake-flask

*Cutaneotrichosporon oleaginosus* ATCC 20509 was used for genetic manipulation studies and maintained on Yeast extract Peptone Dextrose (YPD) agar plates containing 10 g/L yeast extract, 20 g/L peptone, 20 g/L glucose, 20 g/L agar. The maintained cultures were stored at 4°C for up to a week. *Escherichia coli* Zymo 10B (Zymo Research, Orange, CA) was used for all cloning purposes throughout this study and maintained on Luria-Bertani (LB) agar (10 g/L tryptone, 10 g/L NaCl, 5 g/L yeast extract, 15 g/L agar) with ampicillin (100 μg/ml) at 37°C.

The inoculum was prepared by transferring a single colony of *C. oleaginosus* into 10 mL YPD broth (10 g/L yeast extract, 20 g/L peptone, 20 g/L glucose, 20 g/L) in 50 mL tubes and incubated at 30 °C, 250 rpm for 18 h in a shaking incubator. Wild-type and other built *C. oleaginosus* transformants were cultivated into minimal media consisting of glycerol as carbon source and urea as nitrogen source with set ratios of C/N (g/g) 30, 120, 175, 200, and 300 (Duman-Özdamar et al., 2022). Biotin (0.2 mg/L), thiamine (1 mg/L), threonine (2 mM), serine (2 mM), and aspartate (2 mM) were added into C/N 120 cultivation medium, and only wild-type was tested in these experiments. C/N 100 minimal medium was used in amino acid supplement experiments as a control to maintain the C/N ratio in the non-amino acid-supplemented group. Cultures were incubated at 30 °C, 250 rpm for 96 h in a shaking incubator. Cells were harvested at the end of incubation and centrifuged at 1780 g, 4 °C for 15 min. All experiments were performed in triplicates.

### 2.6. Plasmid construction and preparation for transformation

Nourseothricin acetyltransferase gene (*NAT*) (GenBank: KY858965.1) was selected as a selection marker for transformation studies, combined with the GPD promoter and terminator (Figure S1). Endogenous ATP-citrate lyase (*ACL1)*, acetyl-CoA carboxylase (*ACC)*, threonine synthase (*TS),* and hydroxymethylglutaryl-CoA synthase (*HMGS)* genes were selected based on the annotation (PRJEB73837) (Nijsse et al., 2024) and combined with an endogenous promoter and a terminator (Table 1). All constructs were synthesized by GenScript Biotech (NJ, The US) by using pUC57 mini as a backbone plasmid (Figure S2, Table S2).

**Table 1.**
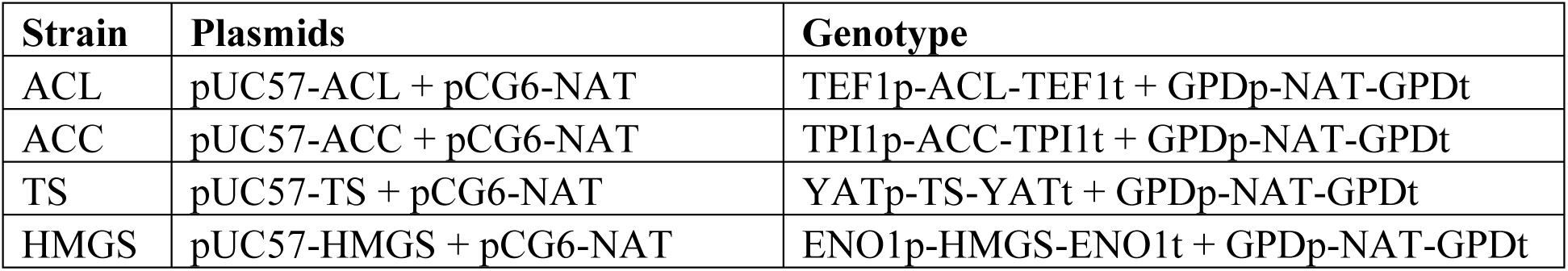
Plasmids and generated *C. oleaginosus* strains.

| Strain | Plasmids | Genotype |
| --- | --- | --- |
| ACL | pUC57-ACL + pCG6-NAT | TEF1p-ACL-TEF1t + GPDp-NAT-GPDt |
| ACC | pUC57-ACC + pCG6-NAT | TPI1p-ACC-TPI1t + GPDp-NAT-GPDt |
| TS | pUC57-TS + pCG6-NAT | YATp-TS-YATt + GPDp-NAT-GPDt |
| HMGS | pUC57-HMGS + pCG6-NAT | ENO1p-HMGS-ENO1t + GPDp-NAT-GPDt |

Synthesized plasmids were transformed into *E. coli* Zymo 10B cells (Cat #T3020; Zymo Research, Irvine, CA, The US) by following the supplier’s instructions, and the transformed strains were stored at −80 °C. *E. coli* cells were grown overnight in 10 ml LB broth with 100µg/ml ampicillin, in 50 mL falcon tubes shaking at 250 rpm at 37 °C. Plasmid DNA was isolated using the GeneJET Plasmid Miniprep Kit (Cat #K0503; Thermo Fisher Scientific, MA, The US). Isolated plasmids were prepared for transformation by linearizing with EcoRI restriction enzyme according to the supplier’s instructions (NEB, Ipswich, MA, The US).

### 2.7. Co-transformation and selection for transformants

Electrocompetent *C. oleaginosus* cells were prepared by transferring 1 mL of overnight culture into 50 mL YPD medium in a 100 mL shake flask and incubated at 30 °C, 250 rpm. When optical density at 600 nm (OD_600_) reached 0.8 to 1.0, cells were harvested by centrifugation at 1780 g, for 15 minutes at 4 °C. The medium was discarded and the pellet was resuspended in 10 mL sterile cold 50 mM sodium phosphate buffer pH 7.5 containing 25 mM dithiothreitol. This solution was incubated for 12 minutes at 37 °C. After incubation cells were harvested at 1780 g for 15 minutes at 4 °C. The pellet was resuspended in 10 mL sterile cold 10 mM Tris/HCl pH 7.6 containing 270 mM sucrose, and 1 mM MgCl_2_ (sucrose solution), and washed twice. Ultimately, cells were resuspended in the pellet in 0.5 mL sucrose solution.

Co-transformation of linearized vectors containing the gene of interest and the nourseothricin (*NAT*) gene was performed by adding approximately 1 µg per vector DNA into 50 µl electrocompetent *C. oleaginosus* cells and incubating them on ice for 5 minutes. Electroporation was performed using a pulse of 0.8 kVolt, 1000 Ohm, 25 µFarad (Bio-Rad, CA, The US) in 2 mm electroporation cuvettes. One mL of YPD broth was added immediately after pulsing. The cells were transferred to a 2 mL Eppendorf tube, and incubated for 2.5 hours at 30 °C, gently mixing the cells every 30 minutes by inversion. 100 µL cells were spread onto YPD agar plate containing 50 µg/mL nourseothricin for primary selection. The negative control was plated on YPD agar. Incubation of the plates was done at 30 °C for 48 hours. Grown colonies were randomly selected and streaked into a YPD agar plate including 100 µg/mL nourseothricin for secondary selection. Transformants were confirmed via colony PCR. The primers in Table S3 and DreamTaq Green PCR Master Mix (2X) were used by following the instructions from the supplier (Thermo Fisher Scientific, MA, The US, #K1081).

### 2.8. Quantification of expression level by qPCR

Overnight cultures of *C. oleaginosus* were grown on YPD broth. After harvesting cells at 1780 g, 4 °C for 5 mins, RNA isolation was performed by Maxwell® 16 Instrument configured with the LEV High Strength Magnetic Rod and Plunger Bar Adaptor and Maxwell® 16 LEV simplyRNA Purification Kit (PROMEGA, WI, The US) by following the supplier’s instructions. As the final step, RNA was eluted in 35 μL of nuclease-free water. 20 μL of eluted RNA was treated with TURBO DNA-*free*^TM^ Kit (Invitrogen, MA, The US, #AM1907) by using routine DNase treatment protocol provided by the supplier. The concentration and quality of RNA were detected with NanoDrop 2000 (Thermo Fisher Scientific, MA, The US). cDNA was synthesized using High-Capacity RNA-to-cDNA™ Kit (Thermo Fisher Scientific, MA, The US) and stored at −20 °C prior to qPCR experiments.

qPCR reactions were prepared by using cDNA, designed primers (Table S3), and PowerTrack™ SYBR Green Master Mix (Applied Biosystems™, MA, US, #A46112) based on instructions from the supplier. QuantStudio™ Design & Analysis software (v1.5.2) was used for experimental design and relative quantification calculations with QuantStudio 1 qPCR equipment (Applied Biosystems™, MA, The US).

### 2.9. Analytical methods

The growth of oleaginous yeasts was monitored every 24 h by measuring the OD_600_. Measured absorbance was converted into dry cell weight for *C. oleaginosus* as explained by Duman-Özdamar et al., 2022. The total fatty acids were determined quantitatively with a gas chromatograph (GC), 7830B GC systems (Aligent, Santa Clara, CA, The US) equipped with a Supelco Nukol^TM^ 25357 column (30m x 530 µm x 1.0 µm; Sigma-Aldrich, St. Louis, MO, The US), hydrogen as a carrier gas. Samples were initially prepared as described by (Duman-Özdamar et al., 2022). Chloroform was evaporated under nitrogen gas and lipid in the tubes was dissolved in hexane before GC analysis.

### 2.10. Regression models

All computational analysis was performed with R version 4.0.2 (R Core Team, 2020). The relationship between the responses (*Y*) and factor (*x*) was expressed by fitting a second-order polynomial equation: 𝑌 = 𝑎𝑥^2^ + 𝑏𝑥 + 𝑐. The quality of the regression equations was assessed according to the coefficient of determination (R^2^). Statistical analysis of the model was performed using Analysis of Variance (ANOVA) and p < 0.05 was considered significant. The peak of the quadratic curves was calculated and identified as optimal levels of the C/N ratio.

### 2.11. Data availability

Supplementary files are available at ZENODO https://doi.org/10.5281/zenodo.10839375, supplementary figures and tables are available at the end of the file.

## 3. Results

Figure 1 illustrates the DBTL approach that we followed to improve *C. oleaginosus* as an efficient oil-producer cell factory.

**Figure 1.**
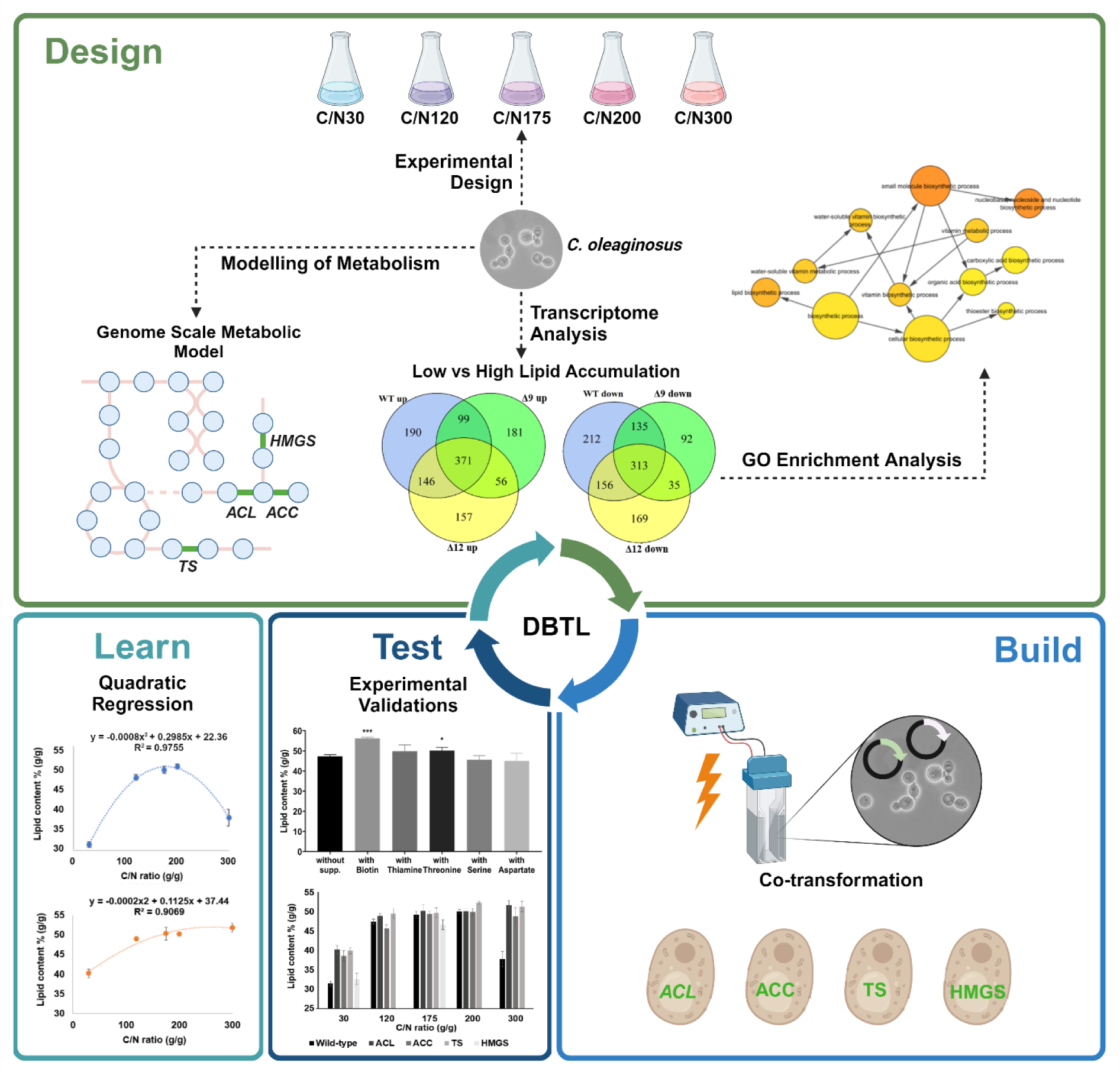
Overview of the DBTL cycle **Design:** We evaluated the transcriptional profile of *C. oleaginosus* by deploying differential expression analysis. Differentially expressed genes were mapped into pathway maps and overrepresented pathways and GO terms were investigated. These investigations were complemented with mathematical modeling of metabolism to identify medium supplements and genetic interventions. Characterization experiments were designed for newly established transformants by selecting the composition (C/N ratio, supplements) of a minimal medium. **Build:** Predicted genetic interventions were implemented by establishing a co-transformation method. **Test:** Predicted medium supplements were tested with wild-type (WT) to validate some of the predicted candidate genes and establish an efficient medium for increased lipid accumulation. Finally, the performance of the built transformants was characterized at various C/N ratios in a minimal medium. **Learn:** Quadratic models were fitted by considering lipid content, dry cell weight, and lipid weight as a response.

### 3.1. Design

In the design step of the DBTL cycle (Figure 1), we evaluated the metabolic capabilities, and selected genes affecting lipid accumulation in *C. oleaginosus* for subsequent modulation, and designed experiments for the characterization of newly built strains.

#### 3.1.1. Differential expression analysis to explore the transcriptional landscape

Three *C. oleaginosus* strains (WT, Δ9, and Δ12), were grown in high C/N and low C/N ratio (10 % CDS, and 60 % CDS) medium, and lipid production was measured (Supplementary File A, Table S1). As expected, higher lipid weights were measured for all strains at 10 % CDS as this provided a more N-limiting environment and promoted higher lipid accumulation. Transcriptomics analysis was performed by collecting samples from these experiments to identify differentially expressed genes in these contrasting conditions (Figure S3). Applying these filters revealed the upregulation of 806 genes in WT, 707 genes in Δ9, and 730 genes in Δ12, and the downregulation of 816 genes in WT, 575 genes in Δ9, and 673 genes in Δ12 (Figure 2). A complete list of DE genes can be found in supplementary file D. Moreover, DE genes in all three comparisons were evaluated, and 371 and 313 genes that jointly up/down-regulated were considered in the following analysis.

**Figure 2.**
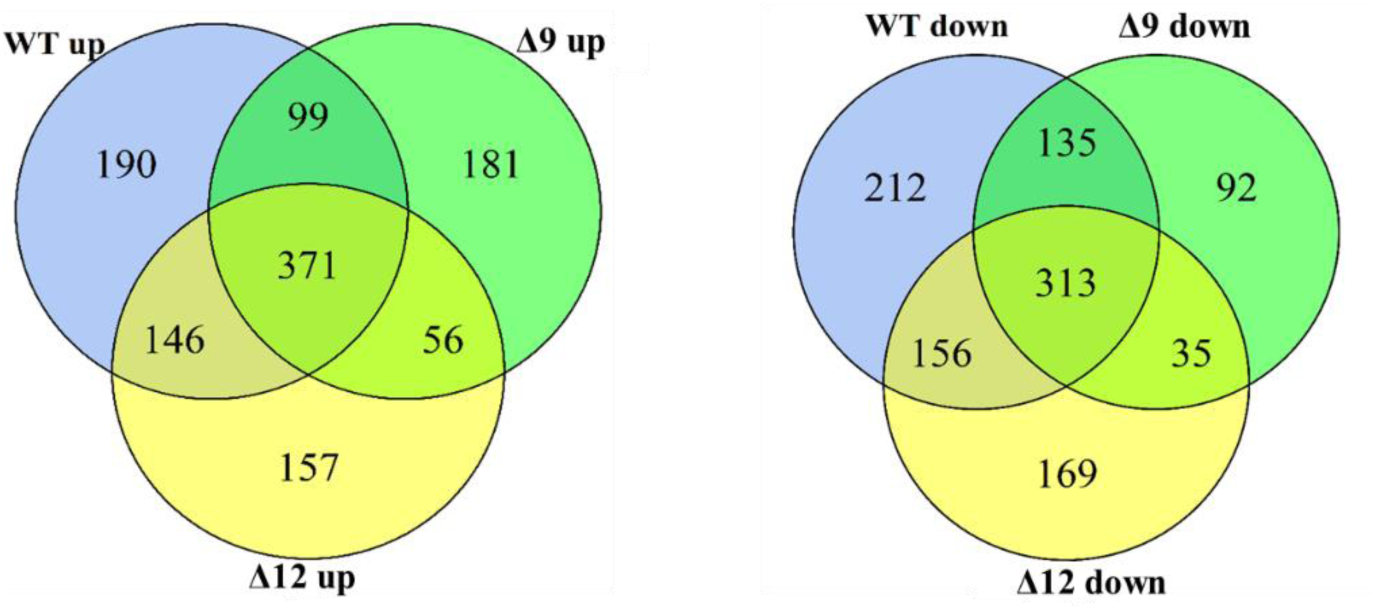
Differentially expressed (DE) genes on WT, Δ9, and Δ12 strains of *C. oleaginosus* on contrasting conditions where strains accumulated high lipid vs low lipid (10% CDS vs 60% CDS). Genes with a higher or lower expression on high lipid accumulating conditions are considered up or down-regulated respectively.

#### 3.1.2. Differentially expressed genes related to metabolism

A metabolic map including glycolysis, citric acid cycle, lipid synthesis, and pentose phosphate pathway was constructed (Supplementary File C). Differences in transcriptional levels in these pathways were represented (Figure 3). Upregulation was observed in the key reactions to initiate lipid synthesis catalyzed by ATP-citrate lyase (*ACL1*), providing citrate, and acetyl-CoA carboxylase (*ACC*), converting citrate to malonyl-CoA. In addition, the whole lipid synthesis pathway and a part of the elongation pathway were upregulated. NADPH is essential for lipid synthesis and a continuous supply is required for lipid accumulation (Ratledge and Wynn, 2002). The NADPH-providing reaction catalyzed by 6-phosphogluconate dehydrogenase was upregulated in higher lipid accumulation conditions. Furthermore, upregulation was observed in the initial step of TAG synthesis catalyzed by glycerol-3-phosphate O-acyltransferase (*GPAT*).

**Figure 3.**
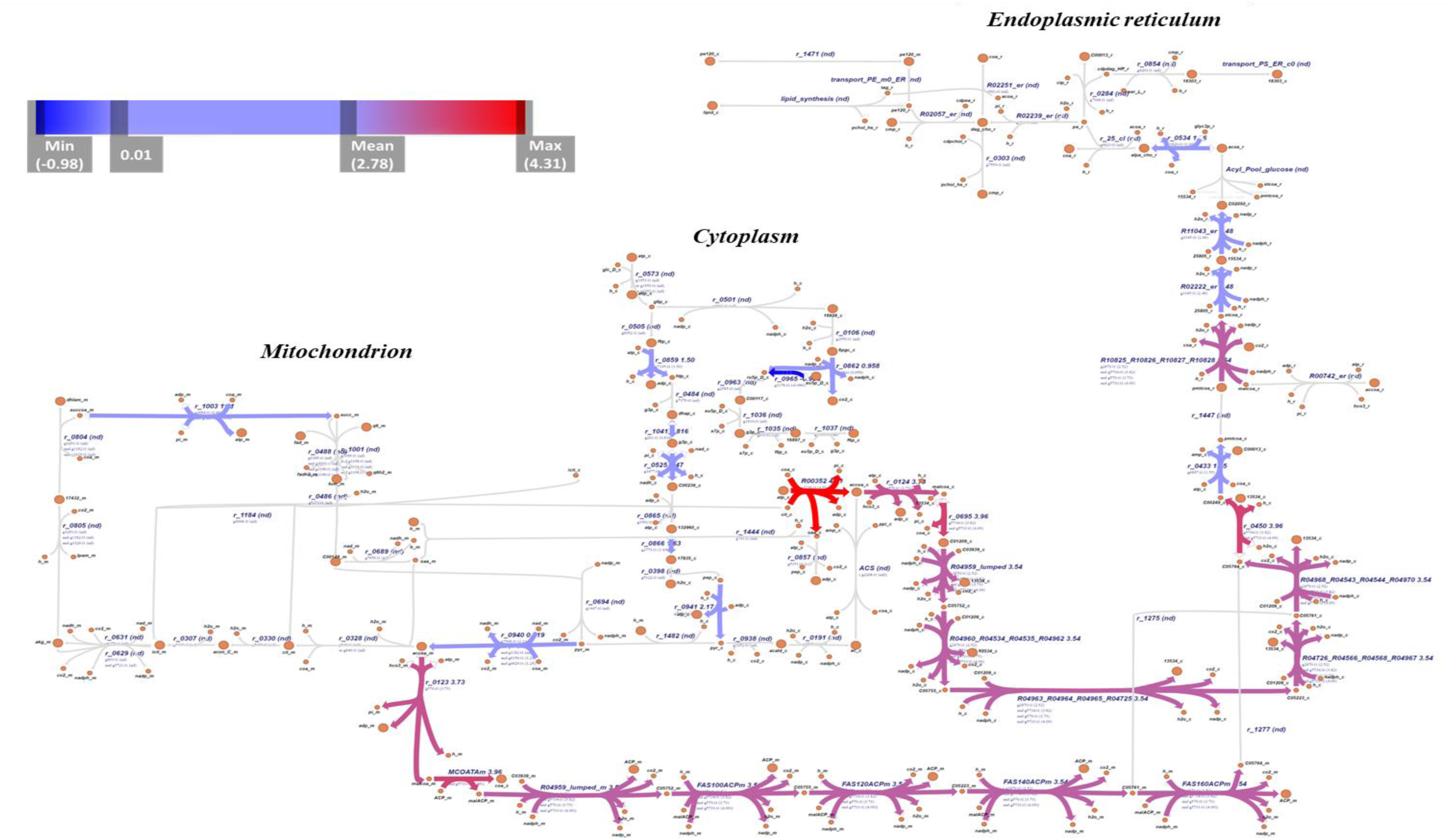
Metabolic map consisting of glycolysis, pentose phosphate pathway, citric acid cycle, and lipid synthesis pathway of *C. oleaginosus* (compartments: cytoplasm, mitochondrion, endoplasmic reticulum). Differentially expressed genes are illustrated based on their log_2_FC scores via color gradient (upregulated genes: from red to light blue, downregulated genes: light blue to navy blue). Graphical representation was created with Escher (https://escher.github.io/) using the Model (*i*NP636_Coleaginosus_ATCC20509_v2) as a reaction database.

#### 3.1.3. Gene Ontology enrichment analysis on DE genes

GO enrichment analysis was performed on differentially expressed genes in all three comparisons. This analysis highlighted enrichments in genes associated with carbohydrate biosynthesis (35 genes), pyruvate metabolism (12 genes), lipid and fatty acid biosynthesis (23 genes), and vitamin biosynthesis process (13 genes) (Table 2, full results can be found in Supplementary File E).

**Table 2.** Selection of enriched gene ontology (GO) terms of upregulated genes for *C. oleaginosus* in higher lipid accumulation condition. (FDR: the false discovery rate, Cluster frequency: the ratio of associated genes in a sample and the total input gene associated with a GO term, Total frequency: the ratio of the number of genes in the genome that are associated with that GO term and the total annotated genome size.) Full results can be found in Supplementary File E.

| GO ID | GO description | FDR | Cluster frequency (171) | Total frequency (3140) |
| --- | --- | --- | --- | --- |
| 16051 | carbohydrate biosynthetic process | 3.09E-14 | 20.4 % | 4.0 % |
| 6090 | pyruvate metabolic process | 6.67E-08 | 7.0 % | 0.7 % |
| 8610 | lipid biosynthetic process | 1.73E-05 | 13.4 % | 4.2 % |
| 6633 | fatty acid biosynthetic process | 4.59E-05 | 4.0 % | 0.4 % |
| 6734 | NADPH metabolic process | 6.89E-05 | 4.6 % | 0.6 % |
| 6767 | water-soluble vitamin metabolic process | 2.02E-04 | 7.6 % | 1.8 % |
| 6766 | vitamin metabolic process | 2.33E-04 | 7.6 % | 1.9 % |
| 6631 | fatty acid metabolic process | 3.10E-04 | 6.4 % | 1.4 % |

Inspection of metabolic functions identified among the overrepresented ontologies highlighted the upregulation of genes associated with providing NADPH, which is required to synthesize fatty acids. Identified genes relate mainly to glycolysis and pentose phosphate pathway glycerol-3-phosphate dehydrogenase (*GUT2*), Glyceraldehyde-3-phosphate dehydrogenase (*GPD1*), 6-phosphofructo-2-kinase (*PFK2*), 6-phosphogluconate dehydrogenase (*GND1*). As expected, lipid and fatty acid synthesis and initiation of fatty acid synthesis were found among the enriched pathways, and key proteins such as fatty acid desaturase, *ACL*, and *ACC* were identified. Additionally, vitamin metabolism and genes annotated biotin and thiamin synthesis were found enriched among the upregulated genes. Therefore, biotin and thiamin additions were selected as media modifications to consider in following DBTL steps.

#### 3.1.4. Comparative Flux Sampling Analysis on GEM

The constrained-based genome scale metabolic model (GEM) of *C. oleaginosus*, *i*NP636_*Coleaginosus*_ATCC20509_v2, was sampled via CFSA by simulating three scenarios, growth, lipid production, and slow growth (Rosmalen et al., 2023). As an output, 1 knockdown and 25 over-expression targets were obtained. Detailed explanation is available at (Rosmalen et al., 2023). Among predicted genetic interventions, ATP diphosphohydrolase which catalyzes the conversion of ATP to AMP was the only downregulation target which indicates the energy requirements of lipid production are higher compared to growth.

As expected, reactions from the fatty acid synthesis pathway, (fatty acyl-acp synthase and stearoyl-CoA desaturase) were suggested as over-expression targets. In addition to the *ACL* and *ACC*, pyruvate dehydrogenase which supplies mitochondrial acetyl-CoA to the TCA cycle, and citrate synthase which produces mitochondrial citrate were listed among the overexpression targets.

Besides lipid synthesis pathway-related targets, unpredicted reactions from the mevalonate pathway such as hydroxymethylglutaryl-CoA synthase (*HMGS*), hydroxymethylglutaryl-CoA reductase, and mevalonate kinase were highlighted. While these reactions involve the consumption of acetyl-CoA, simulations demonstrate a reduction in HMGS flux leads to decreased lipid production. This reflects the relationship between lipid synthesis and mevalonate pathway (Rosmalen et al., 2023).

Furthermore, the model predicted overexpression interventions from amino acid metabolism which are involved in threonine, serine, and aspartate metabolism (threonine synthase (*TS*), aspartate-semialdehyde dehydrogenase, aspartate kinase, glutamine synthase). These predictions were experimentally evaluated in the test step by supplementing threonine, serine, and aspartate into the minimal medium.

Combining all the analyses of the design step led to the identification of four overexpression targets (*ACL1, ACC, TS, HMGS*) (Figure S4) and five medium supplements (biotin, thiamine, threonine, serine, and aspartate) (Table 3), which were transferred to the subsequent build and test steps, respectively.

**Table 3.**
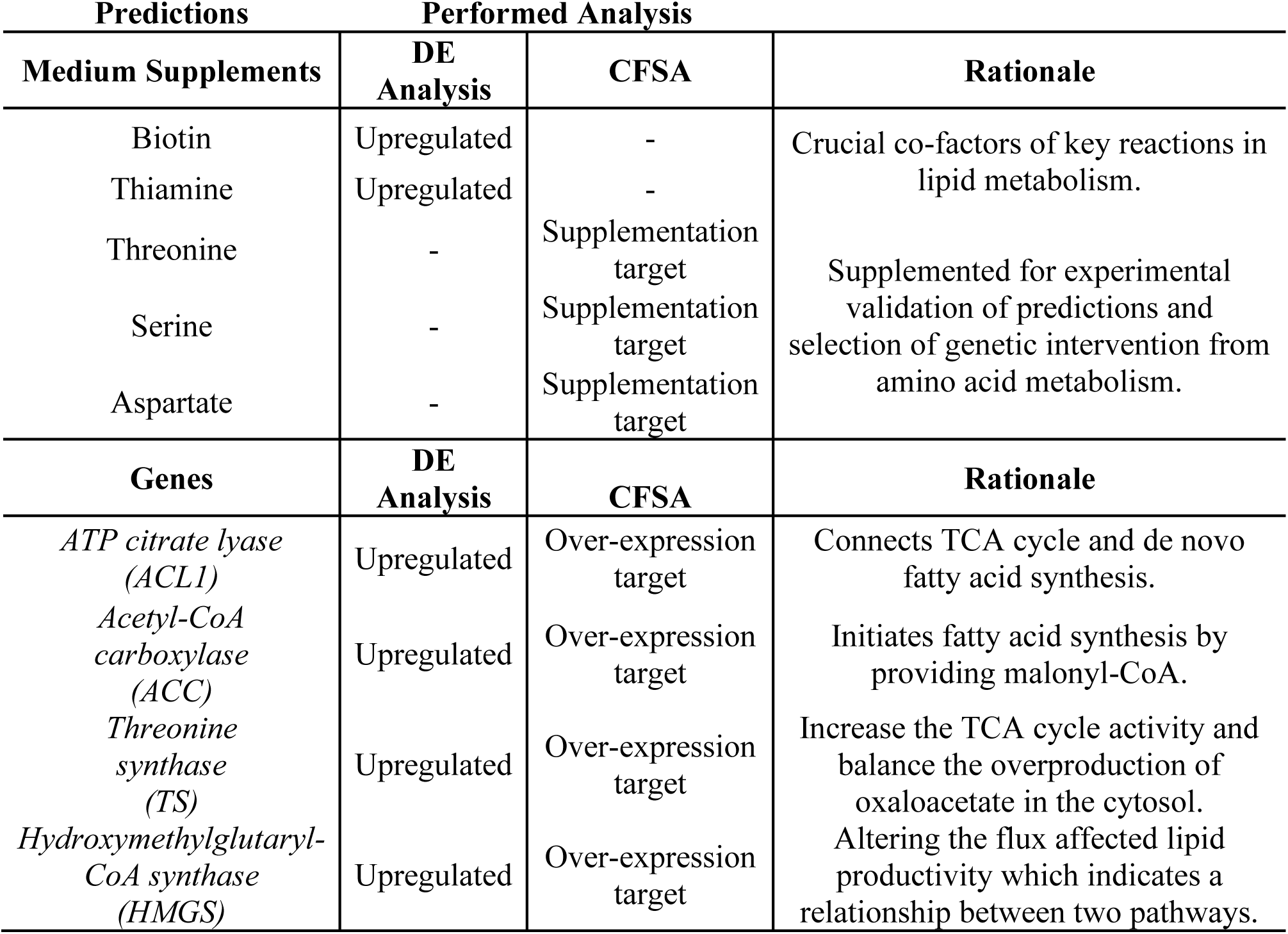
Predicted target genes and medium supplements by investigating transcriptional profile, and analysis on GEM model.

### 3.2. Build

#### 3.2.1. Overexpression of predicted target genes in *C. oleaginosus*

Overexpression of *ACL1, ACC, TS*, and *HMGS* genes (from *C. oleaginosus*) was achieved via the co-transformation of an individual construct for the gene of interest and NAT-containing plasmid. Integration of the gene of interest into the genome of *C. oleaginosus* was confirmed via colony PCR (Figure S5). Afterward, changes in the expression levels were detected by using qPCR for the relative quantification method (Figure 4). Increased expression levels were observed for the recombinant strains, with rather low overexpression for the *TS* gene and very high mRNA levels for *HMGS*. The results confirmed overexpression and thus suitability of the strains for effects on lipid accumulation

**Figure 4.**
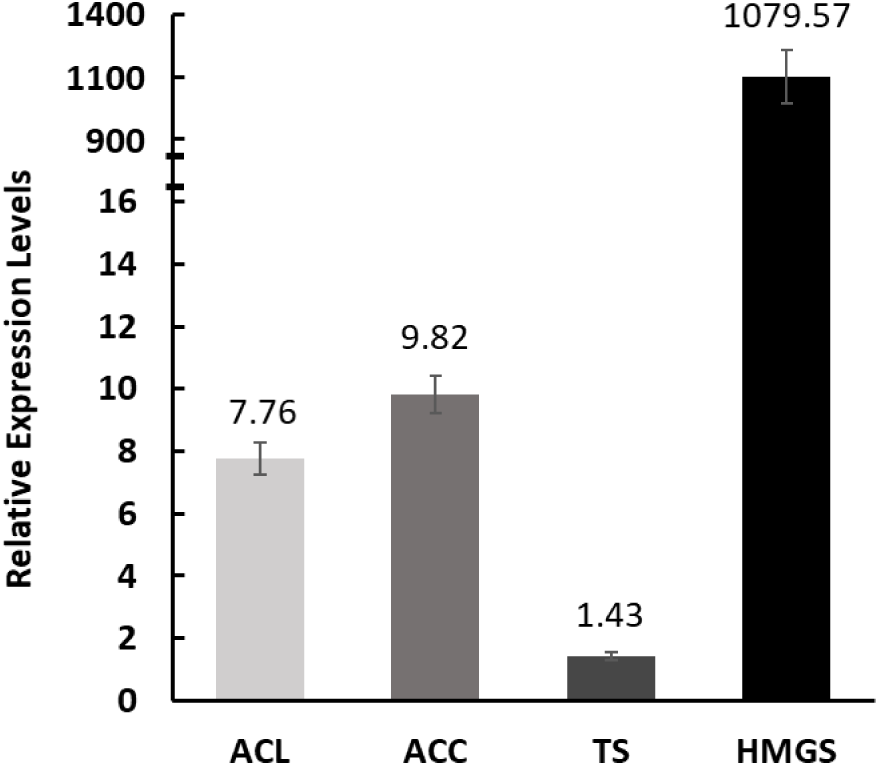
Relative expression levels of *ACL1, ACC, TS,* and *HMGS* in transformants compared to the WT.

### 3.3. Test

In the test step, supplementation predictions obtained on the design step to increase lipid accumulation of *C. oleaginosus* were experimentally tested, and constructed transformants in the build step were characterized at selected C/N ratio cultivation mediums.

#### 3.3.1. Validation of predicted medium supplements

The predictions from the transcriptional profile and GEM model were validated by adding the cofactors (biotin, thiamine) and amino acids (threonine, serine, aspartate) into the cultivation medium at C/N 120. Among these supplements biotin and threonine addition resulted in a significant increase in lipid contents 56.2% (w/w) and 50.2% (w/w) lipid respectively (Table 4). Threonine addition provided around a 17% increase in growth profile. Furthermore, lipid weight was boosted by 31% and 24% when biotin and threonine were supplemented into the medium.

**Table 4.** Lipid content, dry cell weight, and lipid weight of *C. oleaginosus* when biotin, thiamine, threonine, serine, and aspartate were supplemented into a cultivation medium.

| Supplement | Lipid content (% , g/g) | Biomass (g/L) | Lipid weight (g/L) |
| --- | --- | --- | --- |
| No Supplement | 47.27 ± 0.69 | 5.54 ± 0.16 | 2.62 ± 0.11 |
| Biotin | 56.19 ± 0.46 ** | 6.09 ± 0.42 | 3.42 ± 0.26 * |
| Thiamine | 49.81 ± 2.59 | 6.05 ± 0.21 | 3.02 ± 0.26 |
| Threonine | 50.20 ± 1.21 * | 6.45 ± 0.14 ** | 3.24 ± 0.14 ** |
| Serine | 45.65 ± 1.61 | 5.97 ± 0.26 | 2.73 ± 0.2 |
| Aspartate | 48.4 ± 2.94 | 5.68 ± 0.16 | 2.74 ± 0.12 |
| Differences with control (no supplement) were evaluated using a t-test *: $p \leq 0.05$, **: $p \leq 0.01$. | | | |

#### 3.3.2. Characterization of constructed *C. oleaginosus* strains

The effect of overexpressed genes on lipid content, growth, and lipid weight was evaluated by characterizing the strains ACL, ACC and TS in a slightly N-limiting environment (C/N 30), C/N 120, CN 175, which is the optimum C/N ratio determined for WT *C. oleaginosus*, C/N 200 and C/N 300. Mutant overexpressing *HMGS* was tested in C/N 30 and C/N 175 (Duman-Özdamar et al., 2022). Among all tested conditions and constructed strains (Table 5), lipid accumulation varied between 31.5 %, w/w (WT at C/N 30) and 52.35 %, w/w (TS at C/N 200). While ACL, ACC, and TS accumulated higher lipids at C/N 30 and C/N 300 compared to the wild-type, a significant increase occurred for TS also at C/N 200 cultivation medium (Figure 5). Overall, a boost in lipid accumulation was obtained at C/N 300 for all transformants, with ACL lipid content increased by around 40% and lipid weight 1.6-fold upward of WT. Furthermore, overexpression of TS and characterizations at different C/N minimal mediums not only provided an increase in lipid accumulation but also improved the growth profile at three of the selected C/N ratios by up to 55%. As a result of these increases, lipid weight obtained via ACL, ACC, and TS transformants was boosted the utmost 2.5 fold. On the other hand, overexpression of *HMGS* did not alter growth and lipid content.

**Figure 5.**
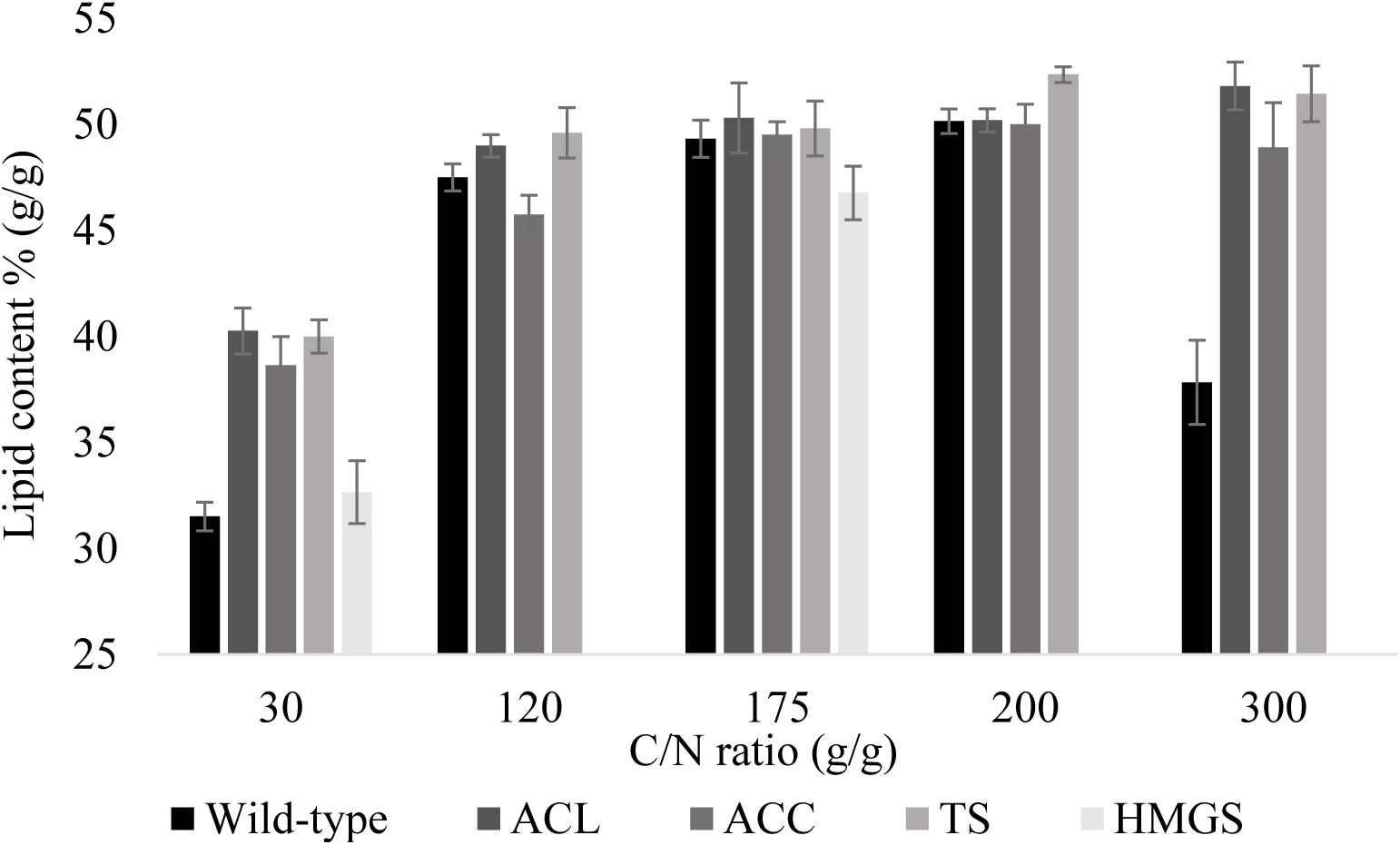
Lipid accumulation profile of WT, ACL, ACC, and TS characterized at C/N 30, 120, 175, 200, 300, and HMGS tested at C/N 30 and 175 (at 96 hours).

**Table 5.** Lipid content, dry cell weight, and lipid weight of WT, ACL, ACC, TS, and HMGS *C. oleaginosus* at various C/N ratio minimal medium.

| C/N 30 Minimal Medium |  |  |  |
| --- | --- | --- | --- |
| Strain | Lipid content (% g/g) | Biomass (g/L) | Lipid weight (g/L) |
| WT | 31.5 ± 0.67 | 4.16 ± 0.23 | 1.31 ± 0.07 |
| ACL | 40.26 ± 1.08 ** | 4.57 ± 0.08 | 1.86 ± 0.10 ** |
| ACC | 38.63 ± 1.37 ** | 4.41 ± 0.16 | 1.70 ± 0.02 ** |
| TS | 40.00 ± 0.77 ** | 5.06 ± 0.13 ** | 2.05 ± 0.08 **<br>* |
| HMGS | 32.64 ± 1.48 | 4.38 ± 0.19 | 1.43 ± 0.12 |
| C/N 120 Minimal Medium |  |  |  |
| Strain | Lipid content (% g/g) | Biomass (g/L) | Lipid weight (g/L) |
| WT | 47.5 ± 0.64 | 5.54 ± 0.16 | 2.63 ± 0.08 |
| ACL | 48.99 ± 0.53 | 5.21 ± 0.08 | 2.55 ± 0.03 |
| ACC | 45.75 ± 0.91 | 5.52 ± 0.29 | 2.52 ± 0.11 |
| TS | 49.61 ± 1.19 | 5.72 ± 0.19 | 2.83 ± 0.08 |
| C/N 175 Minimal Medium |  |  |  |
| Strain | Lipid content (% g/g) | Biomass (g/L) | Lipid weight (g/L) |
| WT | 49.32 ± 0.88 | 5.65 ± 0.12 | 2.79 ± 0.05 |
| ACL | 50.30 ± 1.66 | 5.64 ± 0.08 | 2.83 ± 0.07 |
| ACC | 49.51 ± 0.62 | 5.58 ± 0.08 | 2.76 ± 0.07 |
| TS | 49.81 ± 1.30 | 6.08 ± 0.07 * | 3.03 ± 0.14 * |
| HMGS | 46.77 ± 1.27 | 5.63 ± 0.19 | 2.63 ± 0.08 |
| C/N 200 Minimal Medium |  |  |  |
| Strain | Lipid content (% g/g) | Biomass (g/L) | Lipid weight (g/L) |
| WT | 50.15 ± 0.57 | 6.31 ± 0.13 | 3.16 ± 0.10 |
| ACL | 50.19 ± 0.55 | 6.08 ± 0.05 | 3.05 ± 0.04 |
| ACC | 50.00 ± 0.95 | 6.22 ± 0.35 | 3.11 ± 0.14 |
| TS | 52.35 ± 0.37 * | 6.36 ± 0.08 | 3.33 ± 0.06 |
| C/N 300 Minimal Medium |  |  |  |
| Strain | Lipid content (% g/g) | Biomass (g/L) | Lipid weight (g/L) |
| WT | 37.83 ± 1.99 | 5.33 ± 0.21 | 2.02 ± 0.17 |
| ACL | 51.81 ± 1.13 ** | 6.15 ± 0.22 * | 2.58 ± 0.14 * |
| ACC | 48.91 ± 2.12 ** | 5.96 ± 0.14 * | 2.91 ± 0.07 ** |
| TS | 51.44 ± 1.32 ** | 6.41 ± 0.07 ** | 3.30 ± 0.08 **<br>* |
| Differences with WT (at the same C/N ratio, 96 hours) were evaluated using a t-test *: $p \leq 0.05$, **: $p \leq 0.01$. | | | |

### 3.4. Learn

In the learn step, besides comparing lipid production and growth profile of newly built transformants, the shift at the optimum C/N ratio as a factor for ACL, ACC, and TS on lipid content, dry cell weight, and lipid weight was evaluated by performing quadratic regression analysis (Figure S6). ANOVA was conducted to evaluate the statistical significance and suitability of fitted second-order polynomial equations (Table S4). The quality of the model fit was assessed using the coefficient of determination (R^2^), which is a measure of how much variability in the calculated response values can be explained by the assessed factor. The R^2^ value serves as an indicator of the model’s strength, with closer proximity to 1 indicating enhanced predictive accuracy and model robustness. The R^2^ of models for lipid content, biomass, and lipid content varied between 81.62% (biomass, ACC) and 97.77% (lipid content, WT). All models are regarded as significant as p-values are lower than 0.05.

Based on the selected experimental solution space, the calculated optimum C/N ratio for the total lipid of ACC and TS was shifted towards C/N 250 whereas it was calculated as 185 for WT (Table S5). On the other hand, for ACL, C/N 310 was predicted to obtain higher lipid weight by the regression model. Besides the lipid weight, the predicted optimum C/N for ACL is higher also for lipid content (C/N 257) and biomass (C/N 352) compared to ACC (C/N 239 and 220), TS (C/N 220 and 306) and WT (C/N 179 and 202).

In all, overexpression of ACL, ACC, and TS largely increased the accumulation of fatty acids from slightly N-limiting conditions corresponding to C/N 30 (∼40 % g/g) to a high C/N ratio of 300 (∼52 % g/g).

## 4. Discussion

We deployed the DBTL approach to develop *C. oleaginosus* as a cell factory for improved microbial oil production. We combined transcriptomics analysis and GEM simulations for strain design. We systematically evaluated the changes in the transcription profile and flux distribution of *C. oleaginosus* metabolism in the case of lipid accumulation. The predictions (overexpression targets and medium supplements) were experimentally validated at different stages and growth conditions.

Despite promising results obtained in previous studies for *C. oleaginosus* as a microbial oil producer, studies on developing genetic engineering tools and enlightening the metabolism of this oleaginous yeast are still limited (Bracharz et al., 2017; Chattopadhyay et al., 2021). In this study, we focused on the overexpression of predicted targets and provided a list of efficient constitutive promoters from re-sequenced and re-annotated *C. oleaginosus* ATCC 20509 (Nijsse et al., 2024). Thus, we expanded the repository of the genetic elements (available promoters, endogenous genes, terminators), and established an efficient electroporation-based transformation protocol. Additionally, a co-transformation strategy was established for the first time by using nourseothricin as an antibiotic resistance marker.

Some of the target genes suggested by the CFSA algorithm and the analysis of the transcriptional landscape of *C. oleaginosus* are compatible with genes predicted by other computational analyses and experimentally validated targets for other oleaginous yeasts such as *Y. lipolytica*. In addition, some of the predictions pointed out genes not previously studied in the context of increased lipid accumulation. Reactions catalyzed by *ACL* and *ACC* were predicted as targets by all analysis in the design step. *ACL* has been highlighted as a critical enzyme to connect the TCA cycle and fatty acid synthesis pathway by rerouting citrate to acetyl-CoA and oxaloacetate then *ACC* converts acetyl-CoA into malonyl-CoA and initiates the fatty acid synthesis (Chattopadhyay et al., 2021; Pham et al., 2021). Over-expression of *ACL* and *ACC* has been tested for *Y. lipolytica* and shown that it improved lipid accumulation from 7.3% (w/w) up to around 20% (w/w) when compared to the WT (Ghogare et al., 2020; Tai and Stephanopoulos, 2013; Zhang et al., 2014). However, this increase was observed only when *ACL* from *Mus musculus* was overexpressed in *Y. lipolytica*. Homologous overexpression of *ACL* in *Y. lipolytica* did not show any improvement in lipid accumulation (Blazeck et al., 2014). In this study, we reported for the first time that overexpression of *ACL* improved lipid accumulation from 31.5% (w/w) to 51.8% (w/w). Thus, our results demonstrated that the kinetic properties of *C. oleaginosus ACL* could be superior to that of *Y. lipolytica ACL*. Furthermore, ACC overexpression and biotin addition resulted in an improvement of around 23% in lipid content. This improvement is likely related to the requirement of acetyl-CoA carboxylase of biotin as a cofactor (Magdouli et al., 2020). We believe that optimizing the amount of biotin supplement and the C/N ratio based on the ACC transformant would provide further improvement in lipid accumulation. Thiamine was also predicted as a suitable medium supplement. The addition of this vitamin did not alter the lipid accumulation and growth profile of *C. oleaginosus,* it has been shown that thiamine metabolism has a crucial role in the lipid accumulation of *Y. lipolytica* by characterization of thiamine-auxotrophic strain (Walker et al., 2020). On the other hand, the lipid content of *Y. lipolytica* did not change by supplementing thiamine with a glycerol-based minimal medium (Das et al., 2017).

Simulations on the metabolic model predicted various overexpression targets from the amino acid metabolism related to the synthesis of threonine, serine, and aspartate. (Kerkhoven et al., 2016) reported that in an N-limiting environment, a transcriptomics analysis identified the upregulation of genes related to amino acid metabolism of *Y. lipolytica*. (Wei et al., 2017) modelled the addition of multiple amino acids through *i*_Yali_2.0 GEM model of *Y. lipolytica* which resulted in higher TAG production. Experimental validations resulted in a significant increase in lipid accumulation only when threonine was added to the cultivation medium.

Inspired by these findings and combining them with the output of the CFSA, we overexpressed threonine synthase in *C. oleaginosus* and obtained the utmost 52 % lipid content at C/N 200. Both the addition of threonine and overexpression of TS resulted in higher lipid accumulation in *C. oleaginosus* due to a potential increase in the activity of the TCA cycle providing more citrate to be used for lipid production (Kim et al., 2019). Furthermore, threonine synthesis requires cytosolic oxaloacetate which is produced during citrate conversion to acetyl-CoA by ACL. Therefore, the over-expression of the threonine synthesis pathway could improve lipid production balancing the over-production of oxaloacetate. Additionally, significant improvement was observed in the growth profile with the addition of threonine and for the TS transformant. We believe that this increase is related to the availability of the amino acid source in the cultivation medium and its increased production in vivo.

Another predicted overexpression target was HMGS from the mevalonate pathway. CFSA captured the intricate relationship between the mevalonate pathway and lipid synthesis intrinsic in the used GEM, proposing it as a potential strategy to enhance fatty acid production (Rosmalen et al., 2023). However, overexpression of *HMGS* did not provide improvement in lipid accumulation. While simulations that reduce HMGS flux resulted in diminished lipid production, increasing the flux of this reaction by more than 4 mmol/gDCW/h led reduction in the flux of lipid production (Rosmalen et al., 2023). The relative expression level of HMGS in the transformant was assessed as 1080-fold higher than the WT *C. oleaginosus*. This situation can be related to the basal expression level of *HMGS* in WT. On the other hand, this could be an indication of increasing the expression level of HMGS excessively. We believe that the mevalonate pathway could potentially affect lipid accumulation, and a more detailed analysis of the whole pathway is required to identify suitable genetic engineering strategies.

Lipid accumulation and lipid productivity in addition to the valorization of side streams, bioprocess optimization, and development of low-cost downstream processing have been reported as the main limitations to developing an economically feasible microbial oil production process through oleaginous yeasts (Koutinas et al., 2014; Karamerou and Webb, 2019). Herein we showed that overexpression of *ACL, ACC,* and *TS* resulted in a boosted amount of lipid content, biomass, and lipid weight both at a slightly N-limited cultivation medium and media with elevated C/N ratio. While lipid content and growth of WT altered by increasing the C/N ratio above 200, probably due to substrate inhibition, ACL, ACC, and TS stably accumulated higher amounts of lipids. This highlights the robustness of these strains to variations in substrate concentration. We believe the strains developed in this study have remarkable potential, especially for growing in glycerol-containing side streams i.e. crude glycerol derived from yellow grease or biodiesel production (Bracharz et al., 2017; Cui et al., 2012; Liang et al., 2010).

## 5. Conclusions

In this study, we have deployed an iterative DBTL cycle to systematically establish *C. oleaginosus* as a microbial oil producer cell factory. The analysis of the transcriptional landscape at lipid accumulation conditions *C. oleaginosus* combined with metabolic modelling led to the identification of gene targets and media supplements to increase production.

Predictions were experimentally validated by supplementing biotin, thiamine, threonine, serine, and aspartate into a minimal medium and overexpressing the predicted targets. When biotin was supplemented into a minimal medium we obtained the highest lipid accumulation (56.2% w/w) and lipid accumulation reached 50.2% (w/w) in the case of threonine supplementation.

We have established the first reported electroporation-based co-transformation method for *C. oleaginosus* and by using our annotations we expanded the genetic elements repository for this oleaginous yeast. By considering the limited genetic manipulation tools and protocols, we believe that this advancement will enable more genetic manipulation studies in the future. We have deployed this method to overexpress, four selected target genes (*ACL, ACC, TS, HMGS*).

Characterization of these new transformants was performed at C/N 30 (relatively lower N-limitation), 120, 175, 200, and 300 to evaluate the performance of newly built transformants at various C/N ratio minimal medium. As a result, ACL, ACC, and TS *C. oleaginosus* were superior to the WT at C/N 30 and they revealed robust lipid accumulation up to 52.35 % towards C/N 300 even though the lipid content of the WT dropped to 37.8 %.

Altogether, the established pipeline for efficient strain and medium design with the modeling and data integration efforts has strengthened the basis to use *C. oleaginous* as a platform for microbial oil production, thereby contributing to the development of processes substituting palm oil that are more sustainable.

## Data availability

Supplementary files are available at ZENODO https://doi.org/10.5281/zenodo.10839375, supplementary figures and tables are available at the end of the file.

## Author contributions

All authors conceived and designed the study. ZEDÖ and MSD performed the data analysis. ZEDÖ drafted the manuscript, ZEDÖ, JACV, and EW performed the experiments. VAPMdS, JH, and MSD acquired project funding, and VAPMdS, JH, MSD, and MKJ conceived and supervised the research. All authors reviewed and edited the study. All authors read and approved the final manuscript.

## Acknowledgments

We thank Dirk van Seijst for his valuable contribution to the development of the co-transformation method. We thank Bart Nijsse for his valuable contribution to generating differential expression data. Graphical abstract and Figure 1 were created with BioRender.com.

## Funding

This research was financed by the Dutch Ministry of Agriculture through the “TKI-toeslag” project LWV19221 “Tailor-made microbial oils and fatty acids”.

## Declaration of Competing Interest

JH has interests in NoPalm Ingredients BV and VAPMdS has interests in LifeGlimmer GmbH.

## Supplementary material

**Figure S1.**
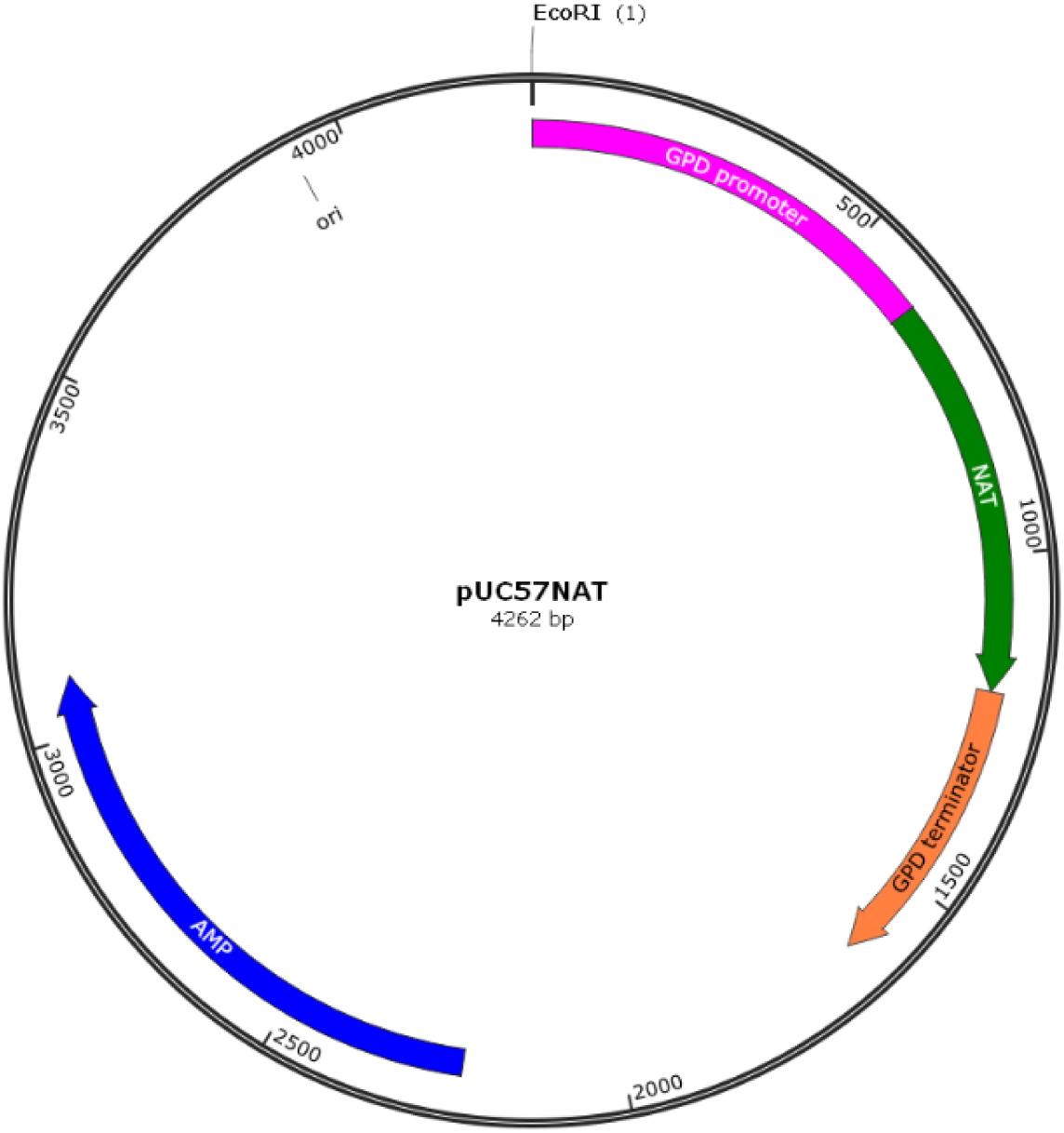
Plasmid map of pUC57NAT containing p*Gpd*, nourseothricin acyltransferase gene and t*Gpd*.

**Figure S2.**
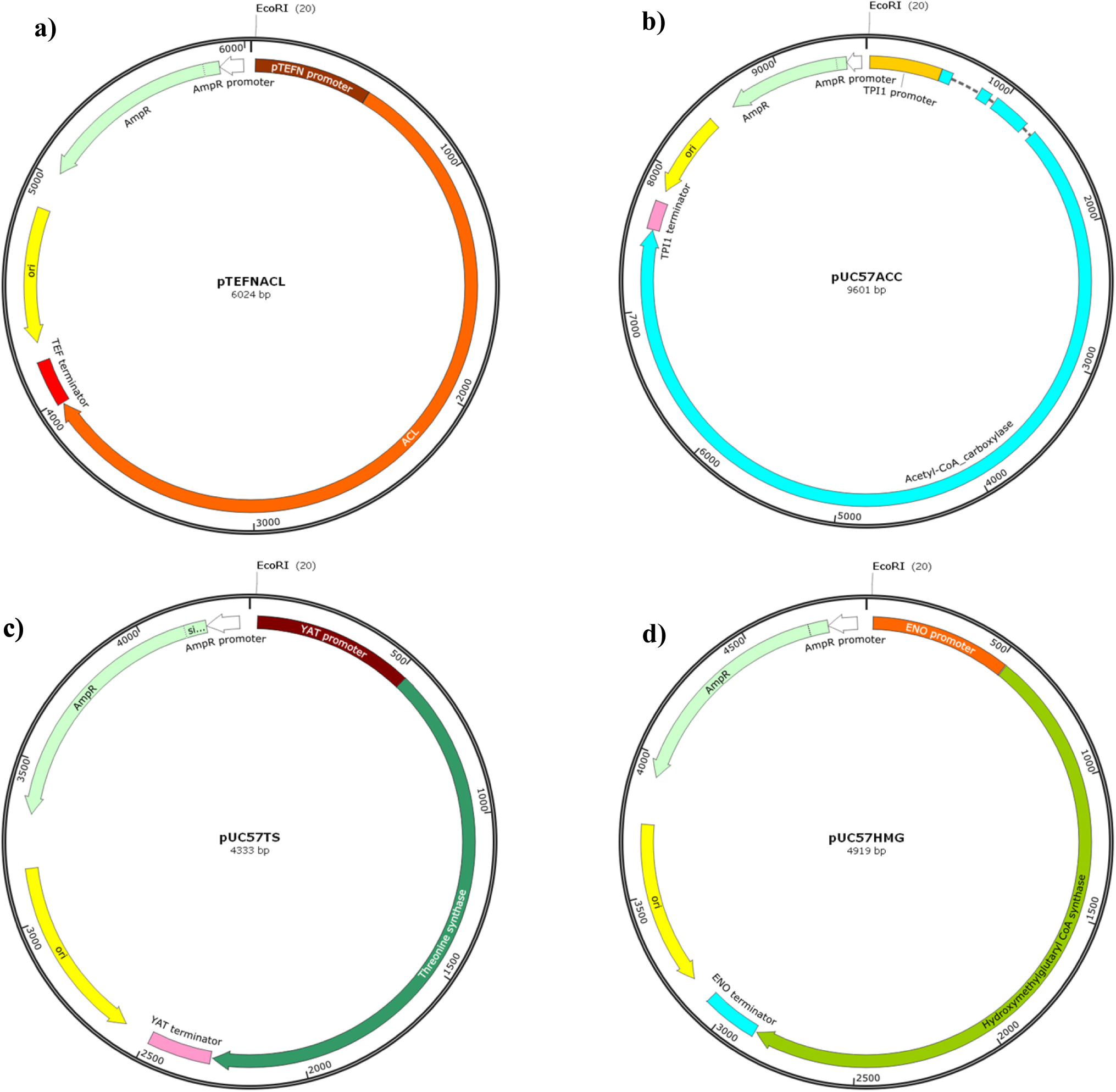
Plasmid maps of overexpression targets containing **a)** TEF1α promoter, ATP-citrate lyase gene, TEF1α terminator, **b)** TPI1 promoter, Acetyl-CoA carboxylase gene, TPI1 **c)** YAT1 promoter, threonine synthase gene, YAT1 terminator and **d)** ENO1 promoter, hydroxymethylglutaryl-CoA synthase gene, ENO1 terminator.

**Table S2.**
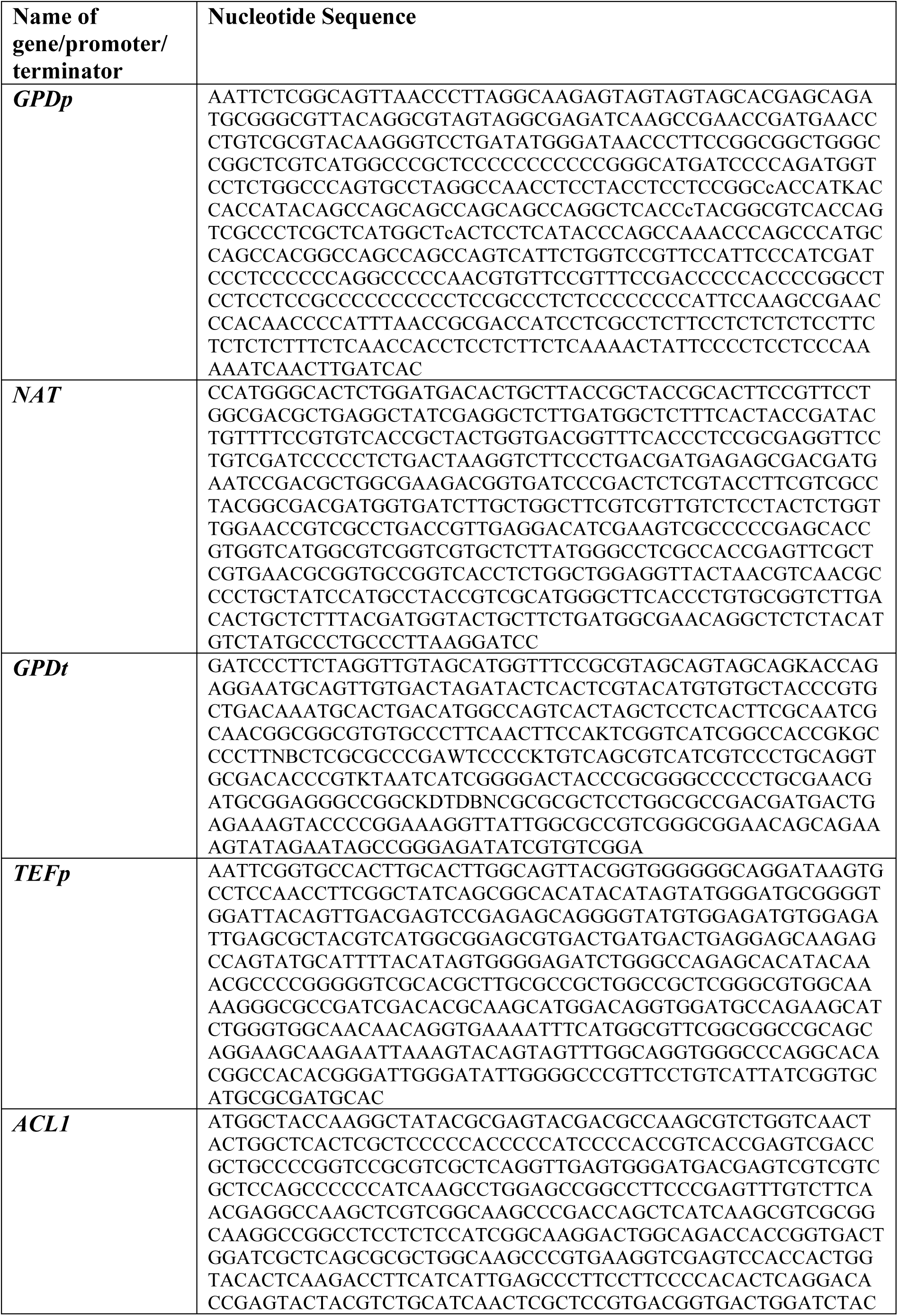
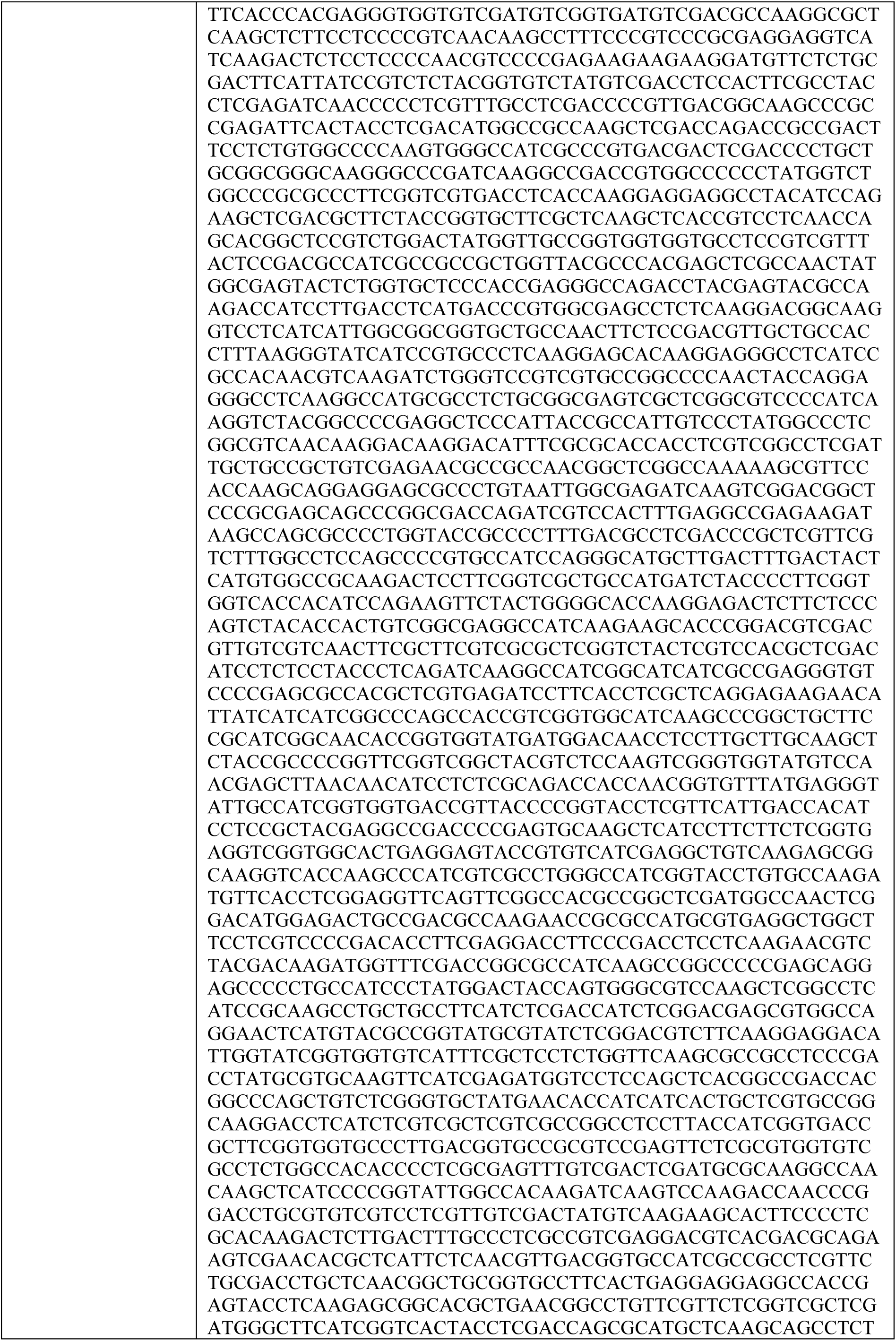
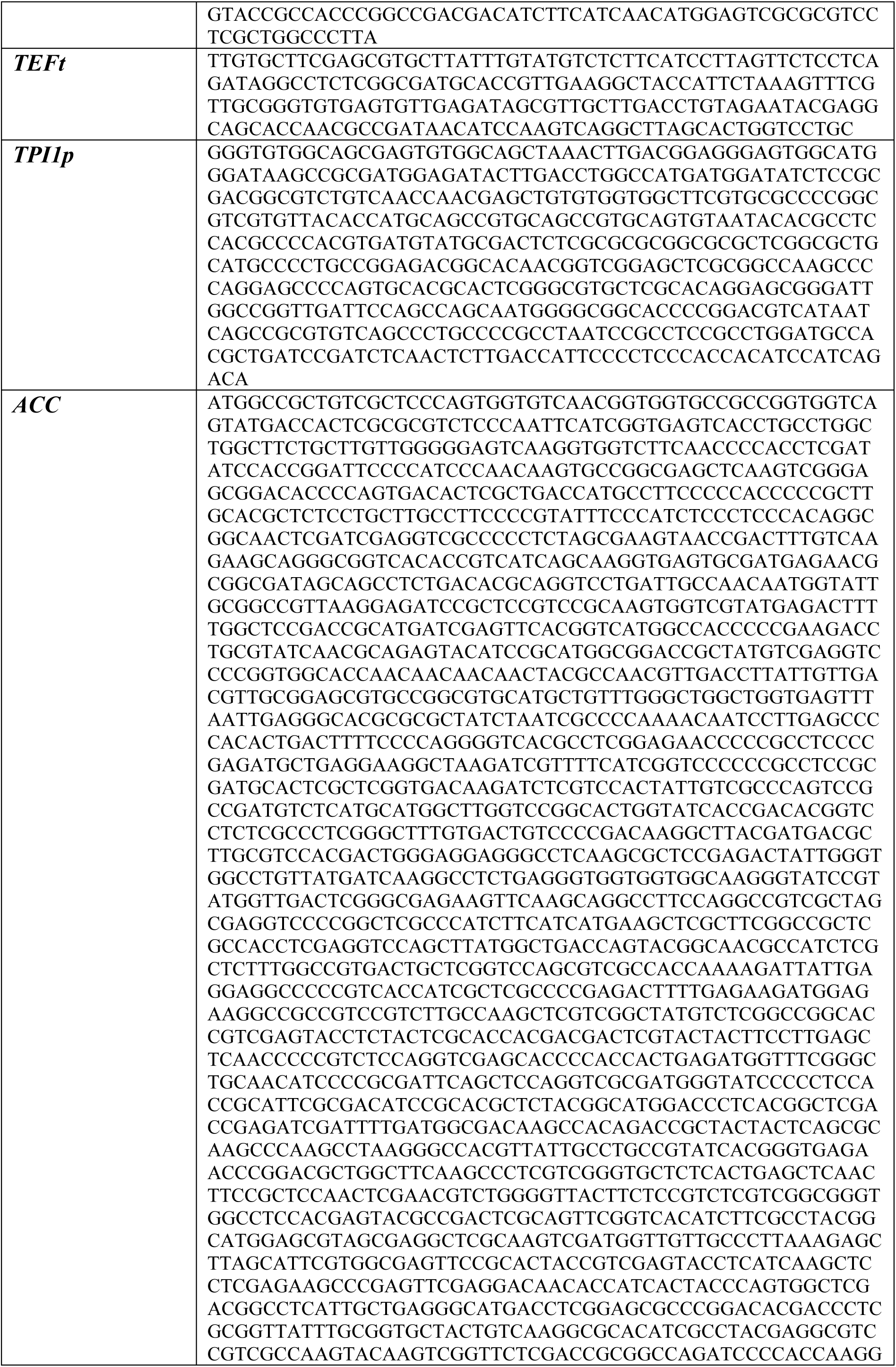
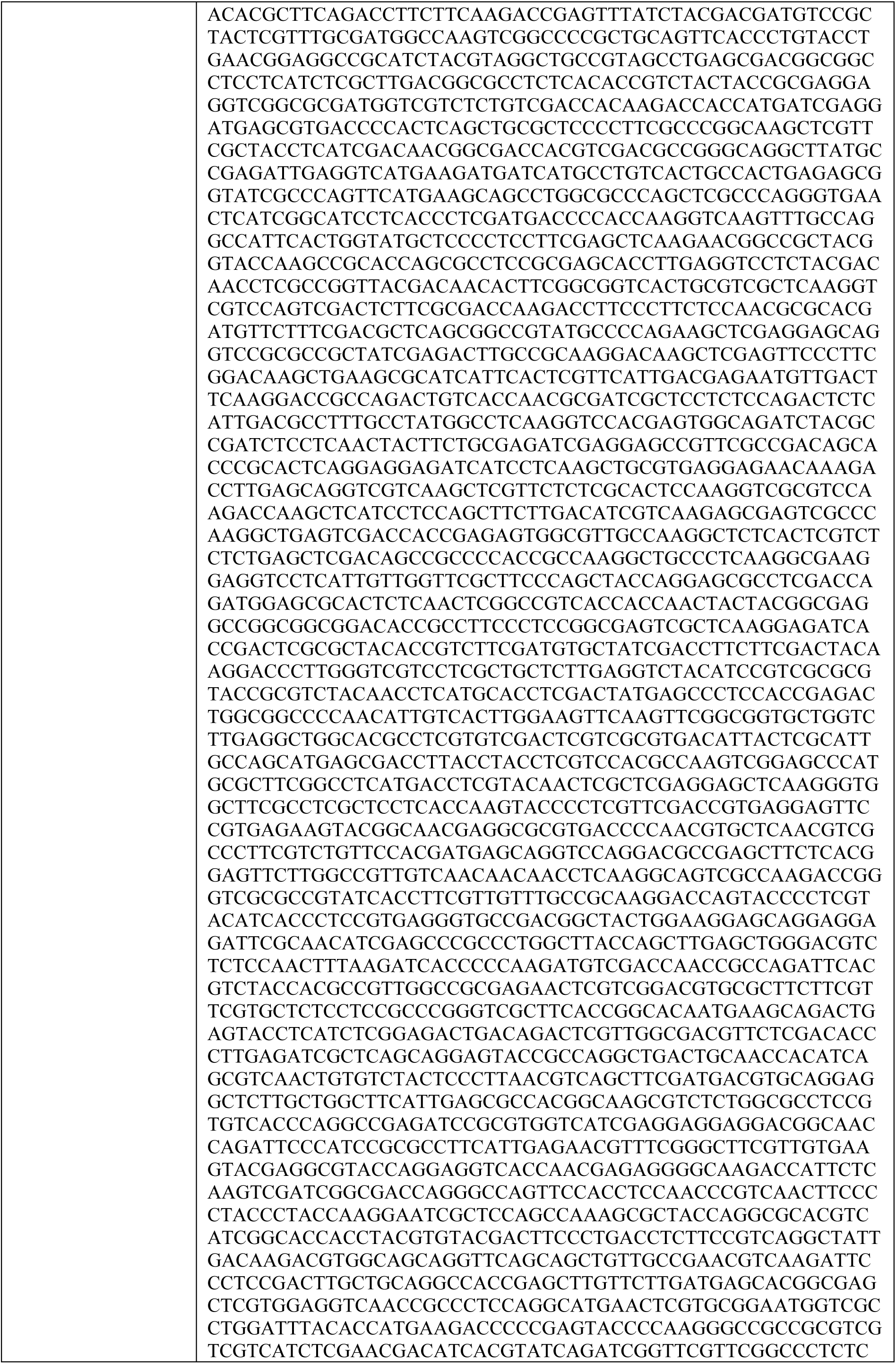
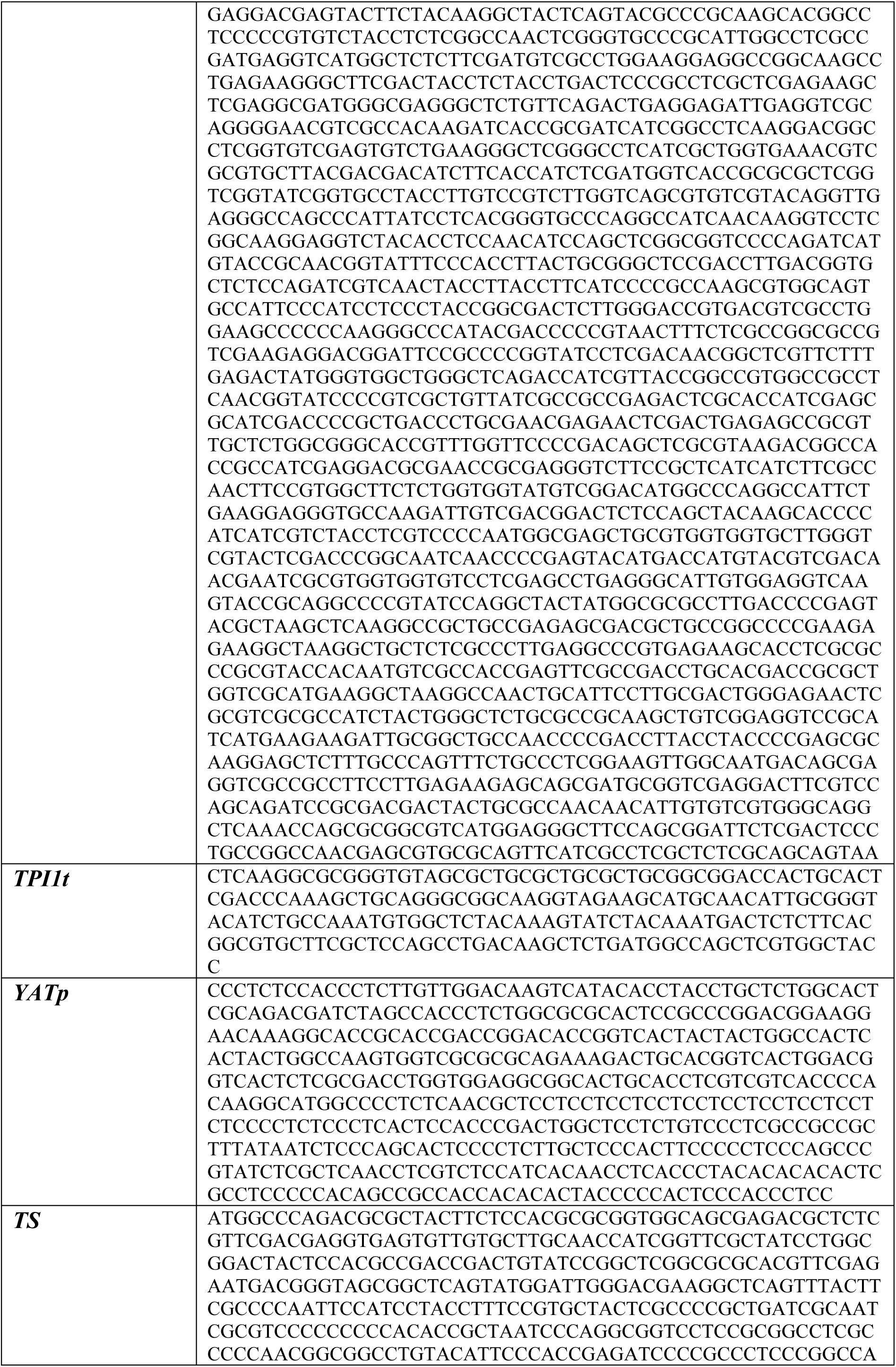
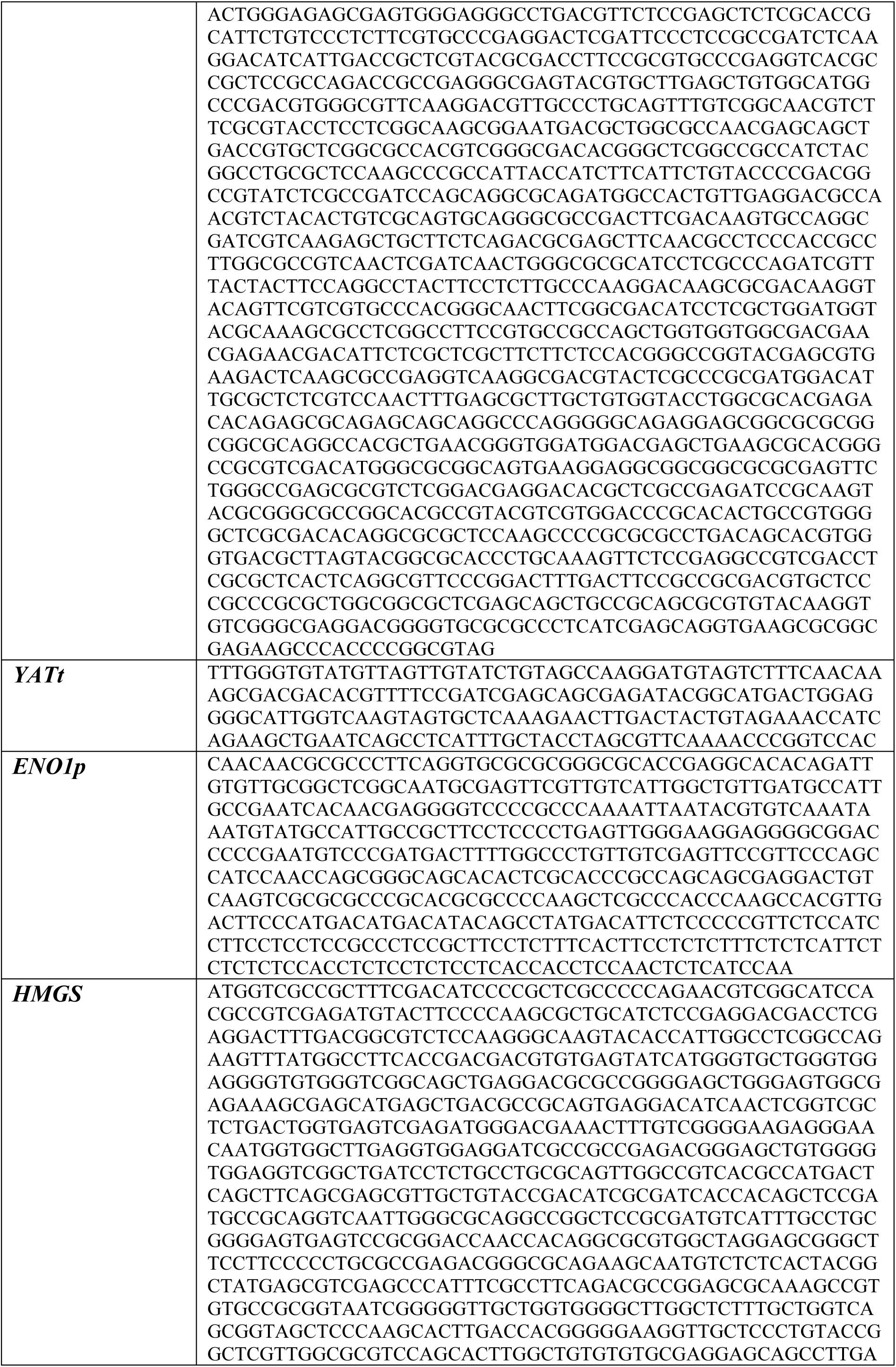
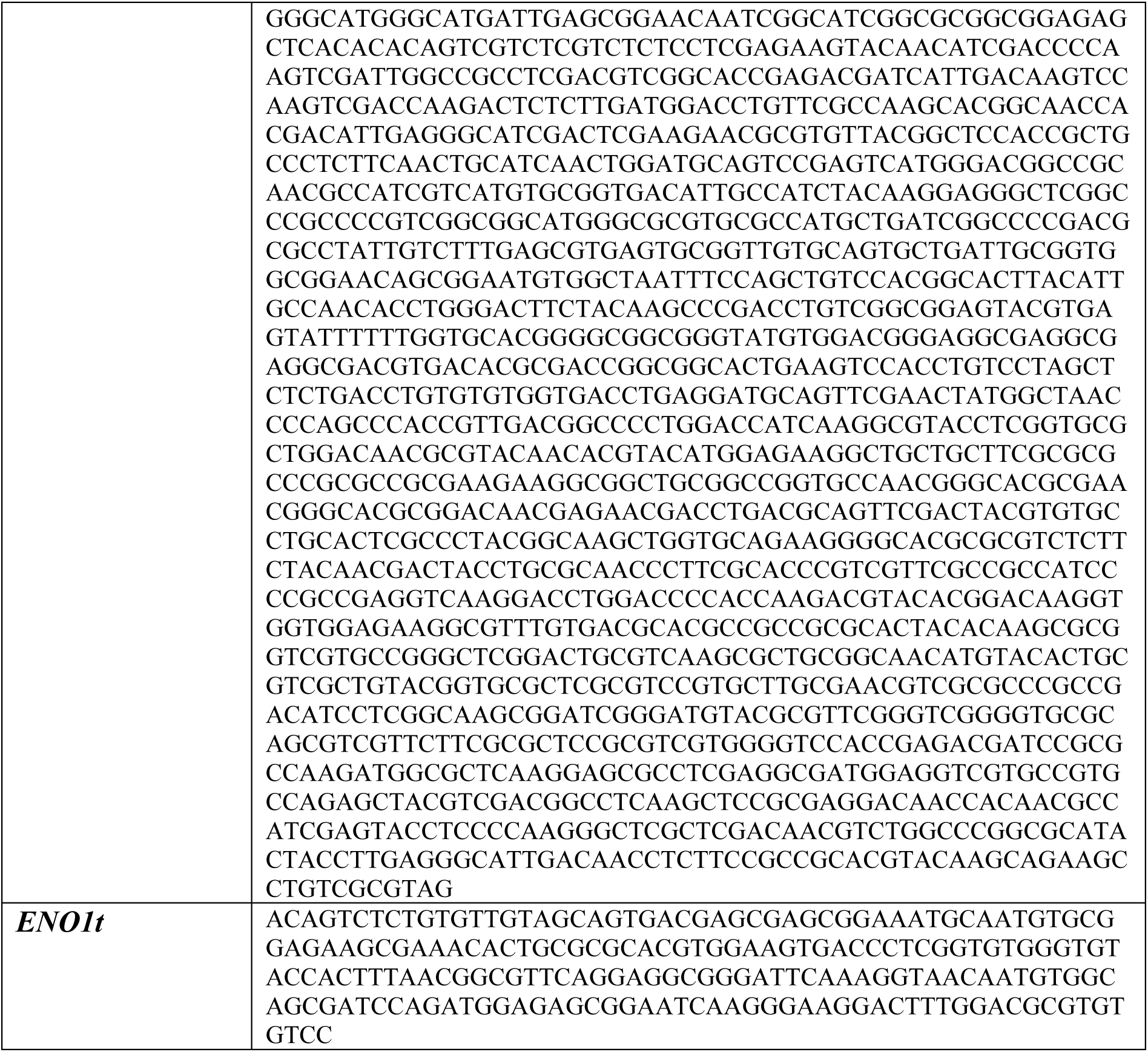
Nucleotide sequences of promoters, genes, and terminators from *C. oleaginosus*.

**Table S3.**
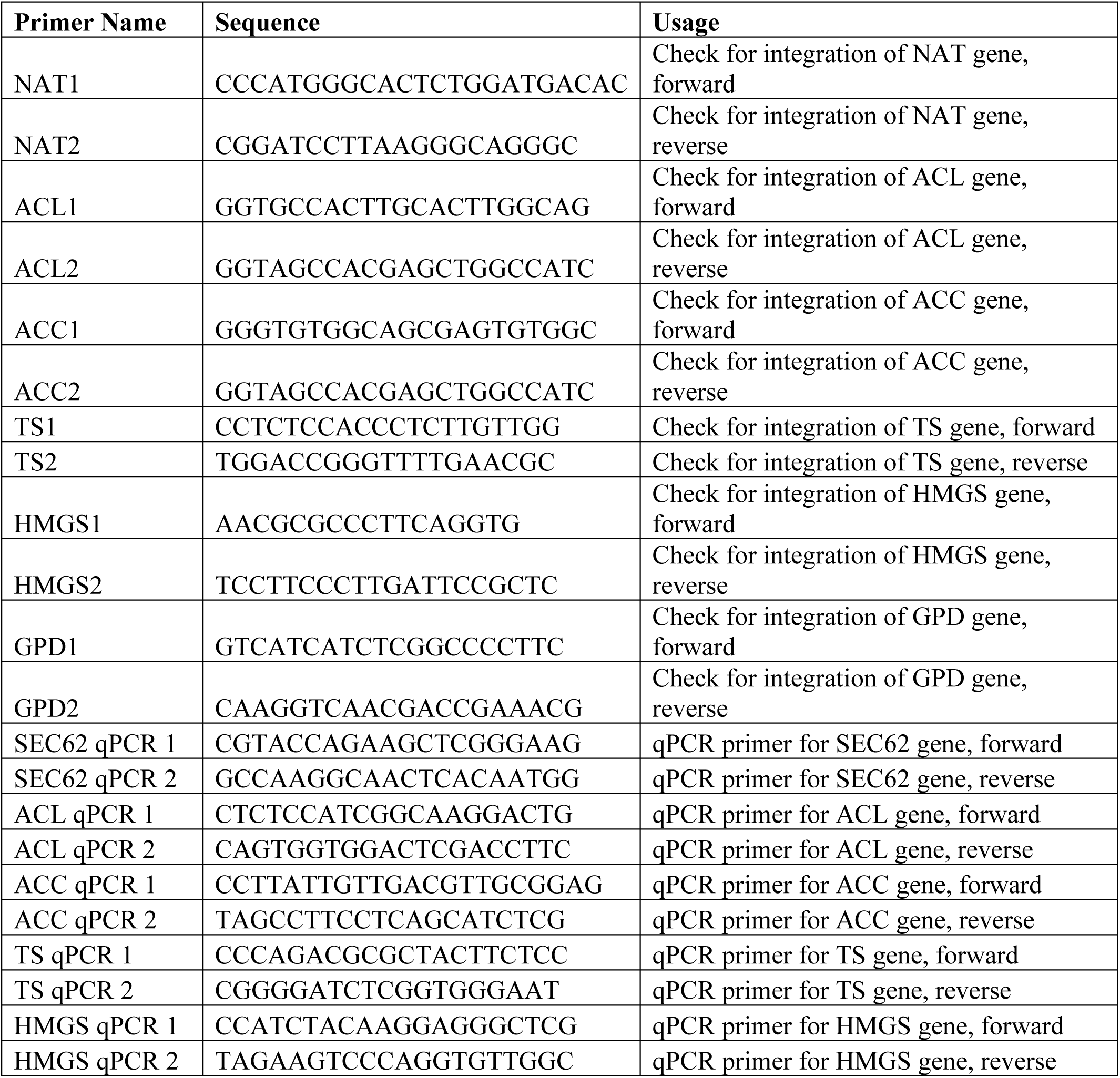
Primers designed for colony PCR and qPCR.

**Figure S3.**
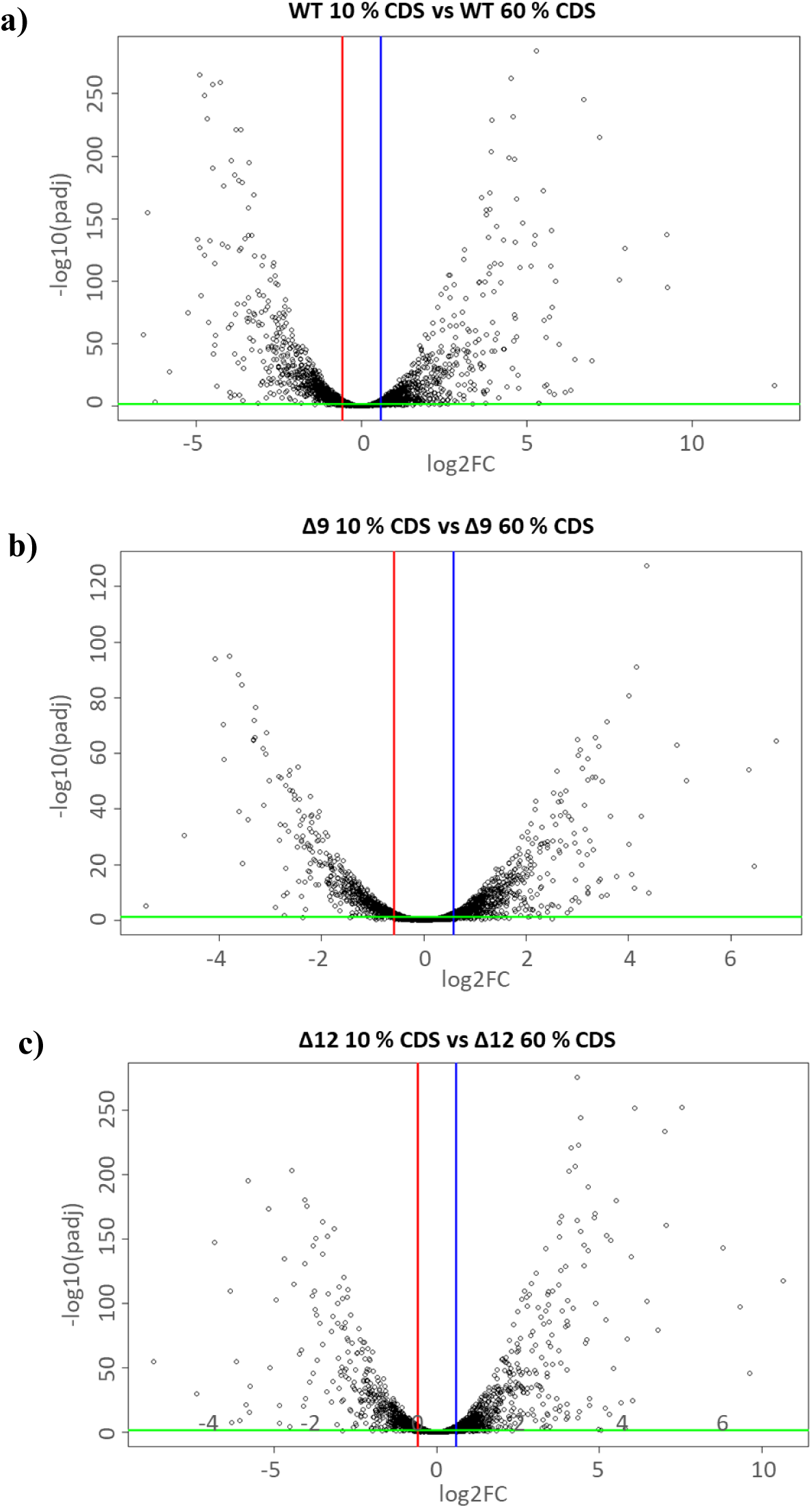
Volcano plots displaying differentially expressed genes and fold change (log2) in expression levels in **a)** WT in 10 % CDS vs WT 60 % CDS, **b)** Δ9 in 10 % CDS vs Δ9 60 % CDS, **c)** Δ12 10 % vs Δ12 60 % CDS. Genes were considered differentially expressed if FC > 1.5 (upregulated) or FC < 1/1.5 (downregulated) and the false discovery rate (FDR) was lower than 0.05.

**Figure S4.**
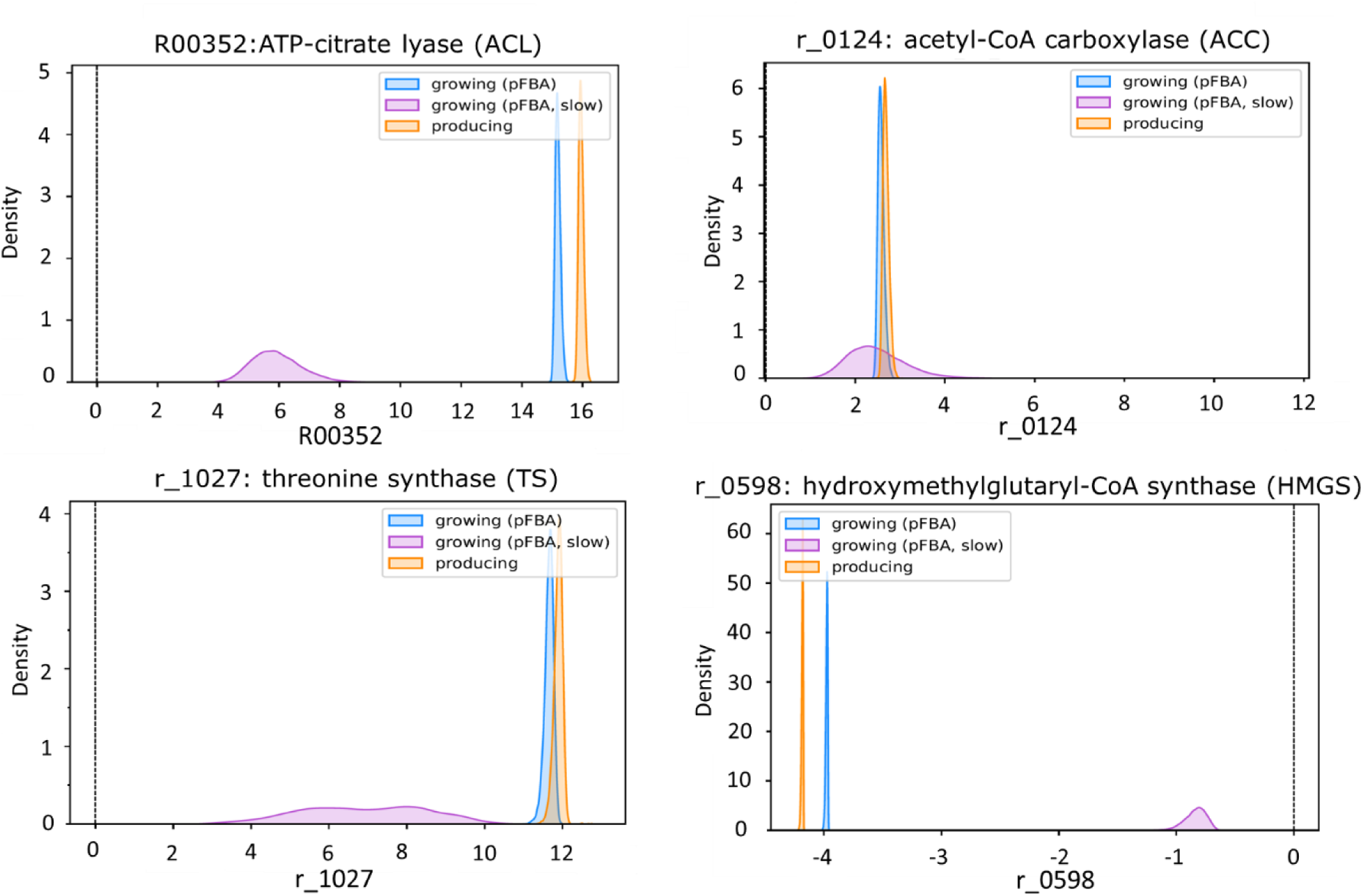
Flux distribution graphs of selected reactions for overexpression in *C. oleaginosus*. CFSA was performed on the metabolic model of *C. oleaginosus* under maximal growth (in blue), maximal production (in yellow), and slow growth (in purple) scenarios.

**Figure S5.**
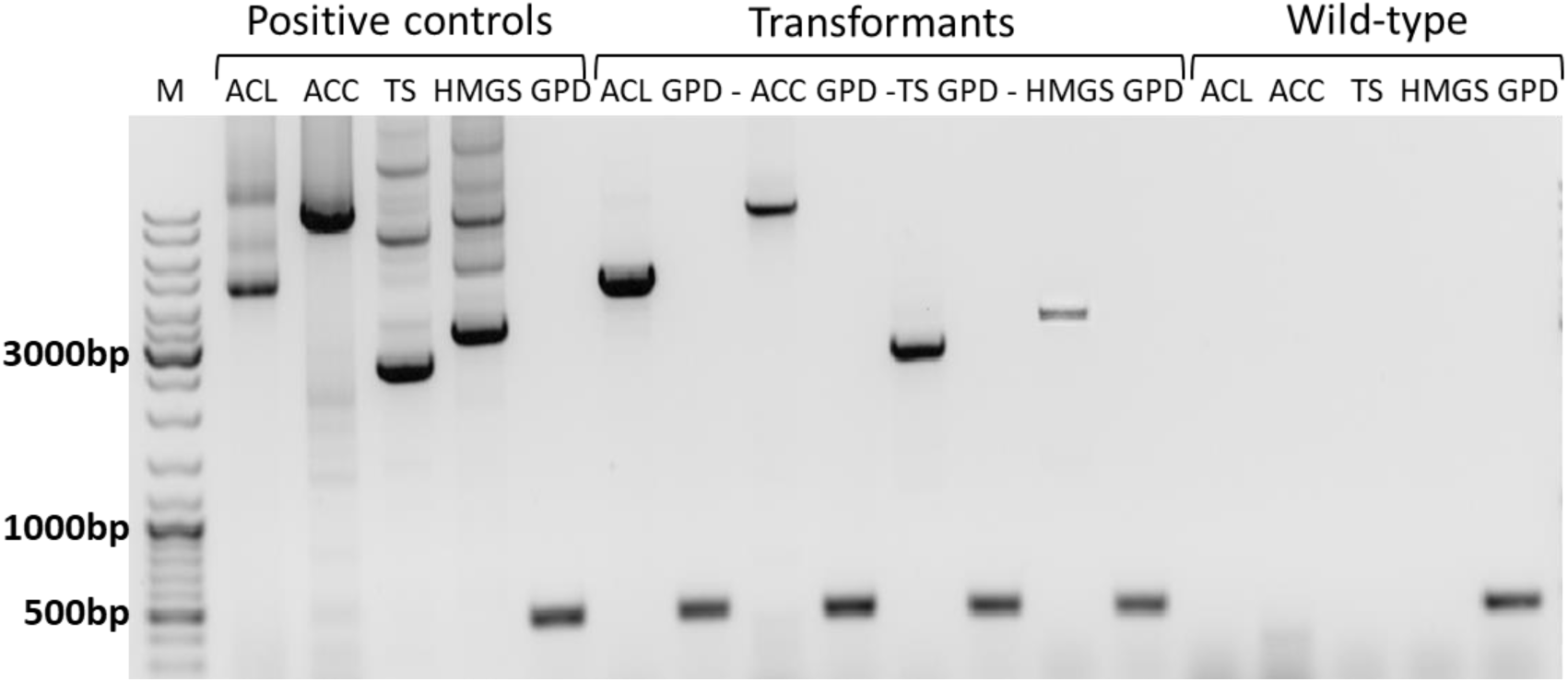
Colony PCR products were run on 1 % agarose gel. Positive controls were prepared by using corresponding primers and plasmids. Quality of gDNA in PCR reaction was assessed by using primers specific for glyceraldehyde-3-phosphate dehydrogenase gene (*GPD*). *ACL*: ATP-citrate lyase gene transformed colony, *ACC*: acetyl-CoA carboxylase gene transformed colony, *TS*: threonine synthase gene transformed colony, *HMGS*: hydroxymethylglutaryl-CoA synthase gene.

**Figure S6.**
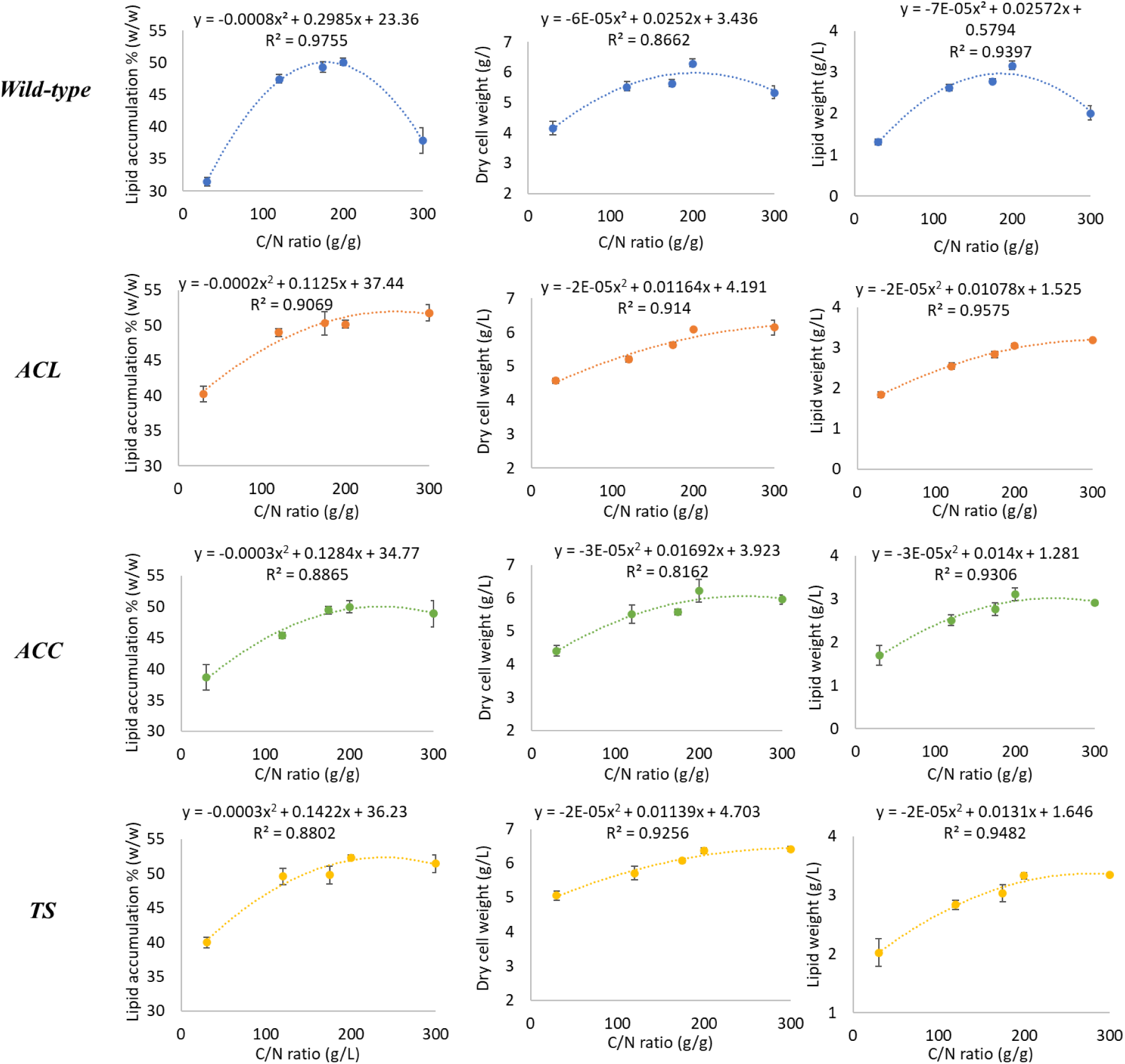
Quadratic regression analysis on lipid accumulation, biomass and lipid content of wild-type, ACL, ACC, and TS *C. oleaginosus* at C/N 30, 120, 175, 200, 300.

**Table S4.**
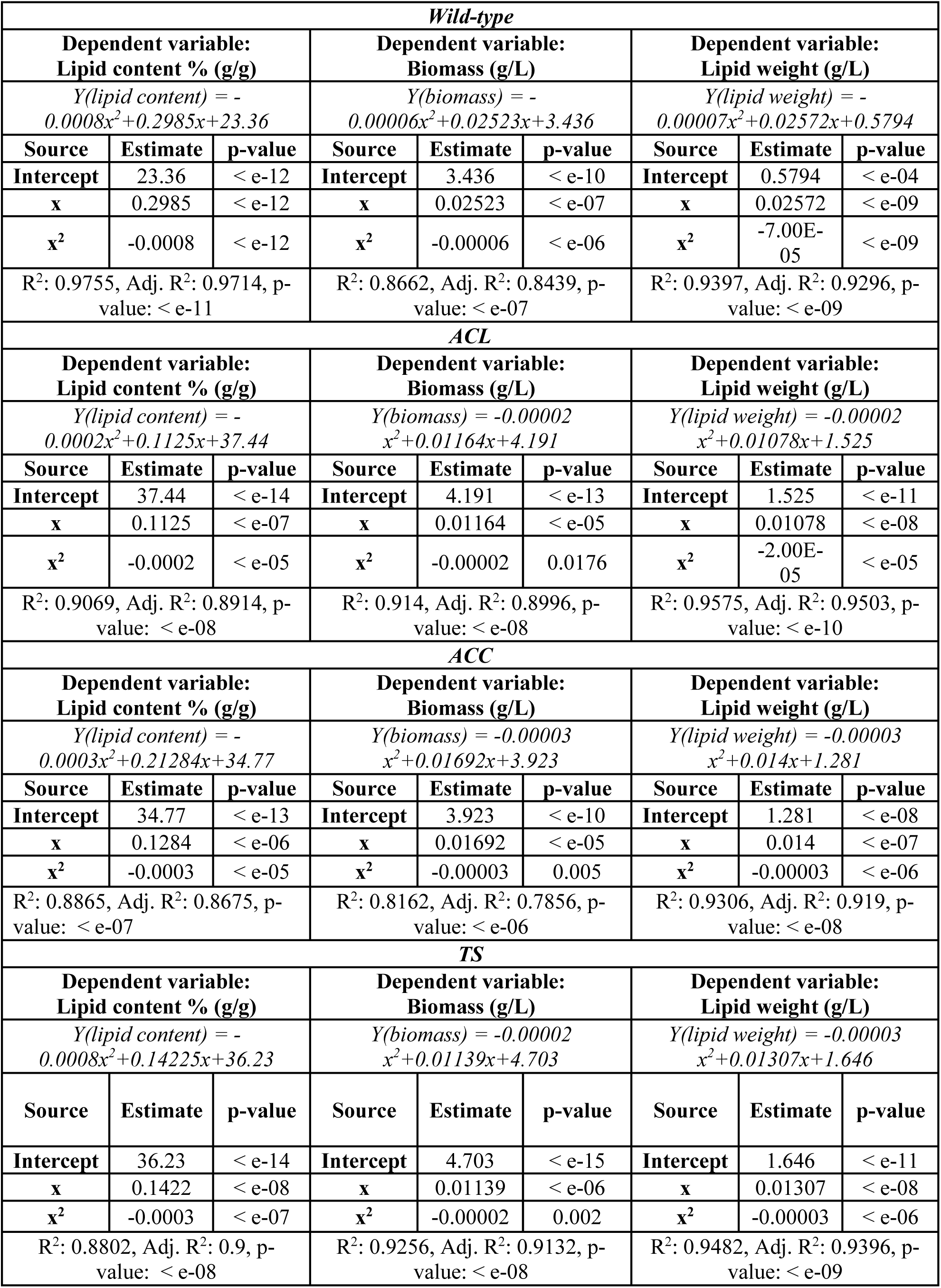
Regression equations, statistics of regression equations for lipid content, biomass, and lipid content of wild-type, ACL, ACC, and TS.

**Table S5.** Calculated optimum C/N ratios and responses (lipid content, biomass, and total lipid) by using built regression models for wild-type, ACL, ACC, and TS.

|  |  | **Lipid content**<br>**% (g/g)** | **Biomass**<br>**(g/L)** | **Total**<br>**lipid (g/L)** |
| --- | --- | --- | --- | --- |
| *Wild-type* | **Predicted optimum C/N (g/g)** | 179 | 202 | 185 |
|  | **Calculated response** | 50.07 | 5.98 | 2.96 |
| *ACL* | **Predicted optimum C/N (g/g)** | 257 | 352 | 310 |
|  | **Calculated response** | 51.91 | 6.24 | 3.20 |
| *ACC* | **Predicted optimum C/N (g/g)** | 239 | 253 | 248 |
|  | **Calculated response** | 50.10 | 6.06 | 3.02 |
| *TS* | **Predicted optimum C/N (g/g)** | 220 | 306 | 250 |
|  | **Calculated response** | 51.89 | 6.44 | 3.28 |

## Notes

https://doi.org/10.5281/zenodo.10839375

